# Trade-offs beget trade-offs: Causal analysis of mammalian population dynamics

**DOI:** 10.1101/2024.08.16.608243

**Authors:** Juraj Bergman, Rasmus Ø. Pedersen, Erick J. Lundgren, Jonas Trepel, Elena A. Pearce, Szymon Czyżewski, Melanie Tietje, Rhys T. Lemoine, Moisès Coll Macià, Mikkel H. Schierup, Jens-Christian Svenning

## Abstract

Survival and reproduction strategies in mammals are determined by trade-offs between life history traits. In turn, the unique configuration of traits that characterize mammalian species give rise to species-specific population dynamics. The dependence of population dynamics on life history has been primarily studied as the relationship between population density and size-related traits. With the recent accumulation of genomic data, the effective population size (*N_e_*) over million-year timescales has become quantifiable for a large proportion of mammal species. Using phylogenetic path analysis, we compared the dependence of population density and *N_e_* on eleven traits that characterize mammalian allometry, diet and reproduction. We found variable impacts of traits on the two metrics of population dynamics across different classifications of mammalian species. For example, we found a negative association between brain size and both population density and *N_e_* overall, but brain size was positively associated with *N_e_* in carnivorans. Dietary specialization had a negative effect on population density, especially in ungulates. *N_e_* of ungulates and primates was strongly affected by time to maturity and weaning age, respectively, highlighting the difference in reproductive strategy between these orders. The relationship between *N_e_* and adult mass showed a gradient in association strength from cold to warm biomes. Together, our findings demonstrate that trade-offs not only characterize life-history evolution, but also extend across different metrics of population dynamics and vary among species groups. This challenges the static nature of the “energetic equivalence” rule and has major implications for selecting appropriate metrics for species conservation and restoration strategies.

## Introduction

The study of life history is central to our understanding of species biology and evolution, as well as informing future conservation and restoration efforts^1–6^. In adapting to a variety of environmental conditions, species have evolved a multitude of strategies emerging from unique configurations of life history traits, with each of these traits allocated a portion of an individual’s energy budget and contributing to its overall fitness^7–11^. The juxtaposition between energy requirements and the benefit of achieving a specific phenotype ultimately results in evolutionary trade-offs between life history traits.

Variation in life history traits gives rise to species-specific population dynamics, characterized by population growth rates^12–16^, fluctuations in population size^17–20^, sex ratios^21–27^ and age structure^28–31^. In mammals, the relationship between population dynamics and traits is most commonly studied by relating population density to allometric traits, such as adult body mass^32–35^. The observed inverse relationship between density and body mass has led to the proposal of the “energetic equivalence” rule (EER; the assumption that the total energy flux within a population is independent of body mass) and scaling laws in ecology^33,36^. Later studies included more complex models of the density-allometry relationship, providing a more nuanced view with phylogenetic relatedness, brain mass and species’ dietary requirements emerging as important predictors of population density^35,37–39^.

However, understanding how phenotypic traits influence population dynamics is complicated by the fact that population density estimates are typically derived from contemporary observations, whereas trait evolution often spans evolutionary timescales. This temporal mismatch is further confounded by extensive anthropogenic impacts on ecosystems in recent times, as well as by the variety of methodologies used to estimate population density, each with its own limitations and potential biases^40^. Consequently, alternative proxies that reflect long-term population dynamics and that can be estimated using standardized approaches are essential for a more complete understanding of the relationship between trait configuration and population dynamics. One such proxy is the effective population size (*N_e_*), now increasingly accessible through advances in whole-genome sequencing and inference approaches^41–44^.

In an idealized setting, *N_e_* measures the rate of genetic drift in a population (random change in the frequency of genetic variants) and determines the efficiency of selection processes^45^. As a measure derived from genetic variation found in natural populations, *N_e_* reflects both demographic and evolutionary processes over long timescales, and is thus expected to provide meaningful insights into how life-history strategies shape long-term population dynamics. For example, larger-bodied species with slower life histories may have smaller *N_e_* due to lower reproductive rates and longer generation times, while dietary specialization can constrain population size and stability, further influencing long-term genetic variation. Offering a long-term evolutionary perspective, *N_e_* estimates may thus complement the ecological insights provided by short-term density estimates. Additionally, recognizing such nuances is critical, especially given the growing use of both density and *N_e_* estimates in macroecological, evolutionary, and conservation studies.

In this study, we modelled population densities and *N_e_*estimates of 380 mammal species as responses to eleven trait variables that characterize species’ allometry, diet and reproduction. We used phylogenetic path analysis (PPA)^46^ to describe trait effects that are common and unique to the two measures of population dynamics, while considering classifications of mammals according to phylogenetic order, trophic level and biome association. Lastly, we tested the plausibility of the “energetic equivalence” rule.

## Materials and methods

### Data compilation

We estimated effective population size (*N_e_*) trajectories of 380 terrestrial mammal species using the pairwise sequentially Markovian coalescent (PSMC) method^41^, following the bioinformatic pipeline from Bergman *et al*. (2023)^47^. This included short read data acquisition from previously published sources, followed by variant discovery using the Genome Analysis Toolkit (GATK)^48^ and inference of the PSMC *N_e_* trajectory. The list of short-read and reference-genome accessions used in the bioinformatic pipeline are reported in Supplementary Table 1 and the pipeline scripts are available at https://github.com/jbergman/mammal_traits (https://doi.org/10.5281/zenodo.18836782).

The phenotypic trait data used as predictors (Supplementary Table 1 and Supplementary Fig. 1) were taken from the COMBINE database^49^, except for the metabolic rate estimates which were taken from Pedersen *et al*. (2023)^50^. We included three allometric traits (adult mass, M; brain mass, B; and metabolic rate, MR) and two dietary traits as defined in González-Suárez *et al.* (2021)^39^ (percentage of diet ^46^made up by animal items, AD; and percentage of diet made up by fruit, nectar and seeds, FS), as predictors. The choice of the two dietary categories is based on the observation that more frugivorous diets are linked to relatively larger brains in primates^51^, while more carnivorous (or folivorous) diets are generally linked to larger body sizes^52^. Additionally, we included six reproductive traits as predictors: time to female maturity, FM; gestation length, GS; weaning age, WA; length of interbirth interval, II; number of offspring per year (calculated as the product of litter size and the number of litters per year), NO; and generation length, GL (Fig. 1). In addition to their relevance to life history strategies of species, these traits were also chosen with the purpose of maximizing the number of species that have been phenotyped for all chosen traits.

**Figure 1.**
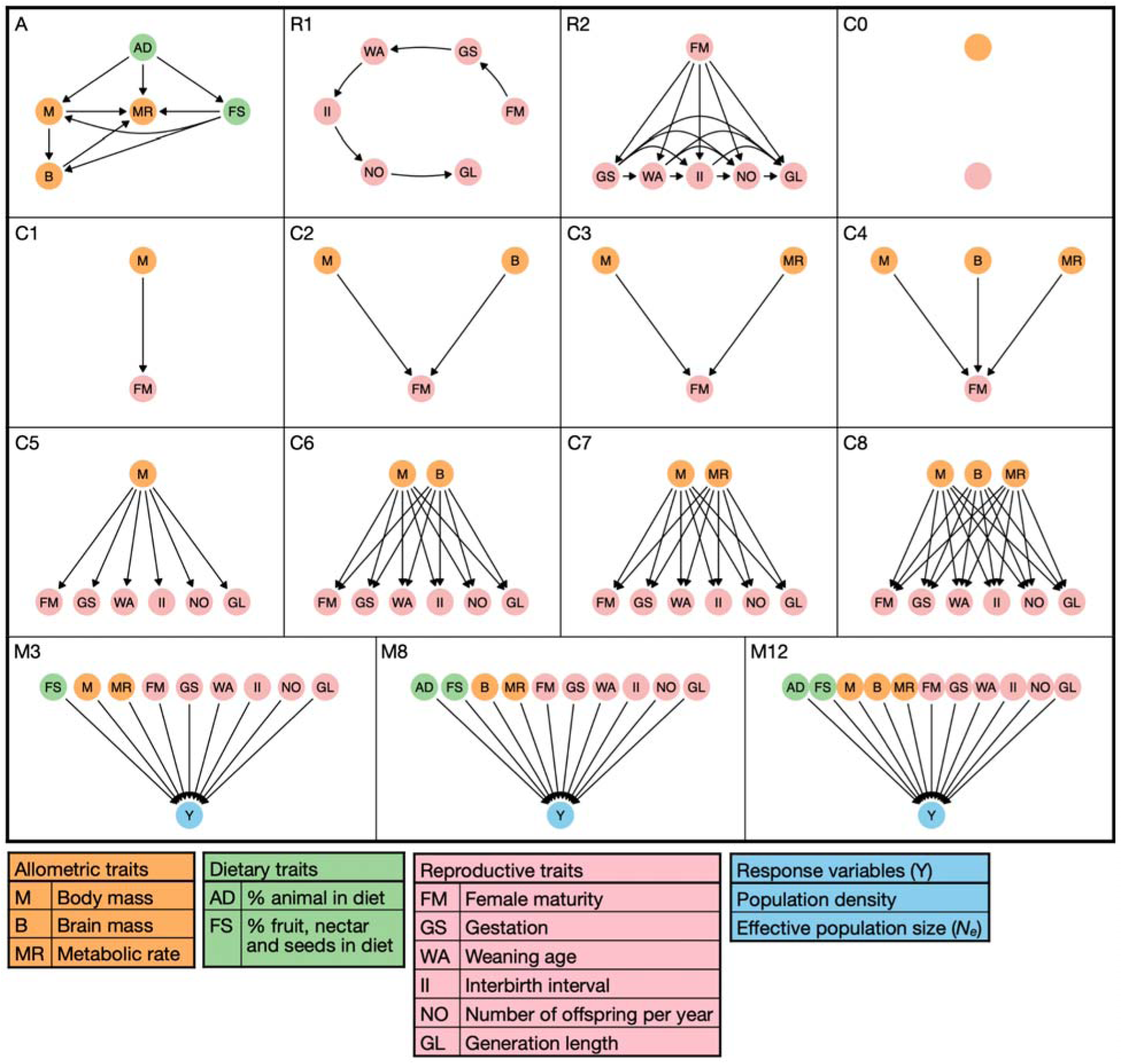
Directed acyclic graphs (DAGs) used in phylogenetic path analysis (PPA) describing causal relationships between phenotypic traits and the response variables. Graph A depicts the relationship between allometric and dietary traits, while graphs R1 and R2 depict the two proposed relationships between reproductive traits. Graphs C0-C8 depict the causal connections between allometric and reproductive traits. Graphs M3, M8 and M12 correspond to relationships between predictor and response variables proposed by González-Suárez et al. (2021)^39^. Given that we considered a single allometry-diet DAG, two reproductive trait DAGs, nine connection DAGs and three predictor-response DAGs, the total number of considered predictor-response networks is 54 (1×2×9×3). All considered predictor-response networks are presented in Supplementary Figs. 3-8.

Two response variables were analyzed (Supplementary Table 1; Supplementary Fig. 2). The first response variable was mammalian population density taken from the Pedersen *et al.* (2023)^50^. These estimates are derived from species’ core areas, body mass and phylogeny and thus represent the best predicted estimates of population densities under low anthropogenic pressure. The second variable we considered was the long-term effective population size of a species (*N_e_*), calculated as the harmonic mean of *N_e_* values from the PSMC trajectory for the period between 100 and 800 kya. This period largely precedes global *Homo sapiens* expansion outside of Africa^53^ and we thus expect minimal anthropogenic impact on *N_e_* estimates during this period for most species and areas^54^. Additionally, ∼800 kya marks the mid-Pleistocene transition (MPT) of glaciation periodicity^55,56^. The chosen period thus coincides with the post-MPT period and encompasses only climate dynamics of 100-ky glaciation cycles. Based on these considerations we argue that the 100-800 kya period can be used as representative of a species’ long-term *N_e_*during the current climate regime.

### Phylogenetic path analysis

We analyzed the relationship between predictor and response variables using phylogenetic path analysis (PPA)^46^. We constructed directed acyclic graphs (DAGs) that represent conditional relationships between variables (Fig. 1). For the DAG describing the relationship between allometry and diet, we adopted the model proposed by González-Suárez *et al.* (2021)^39^ as a base model and added metabolic rate as an additional allometric trait (graph A in Fig.1). Metabolic rate was modeled as a function of all other allometric and dietary traits (as indicated by the directionality of arrows in graph A). For the DAG representing the relationship between reproductive traits, we considered two scenarios. For the first scenario (graph R1 in Fig. 1), we considered a simple model where each reproductive trait is a function of a directly preceding trait within the timeline of reproduction events. The first trait within the reproduction timeline was considered to be time to female maturity, followed by gestation length, weaning age, length of interbirth interval, number of offspring per year, and finally generation length.

The second model of reproduction (graph R2 in Fig. 1) considered each trait within the reproduction timeline to be a function of all preceding traits. For example, gestation length was modeled as a function of time to female maturity (GS ∼ FM), while generation length was modeled as a function of all other reproductive traits (GL ∼ FM + GS + WA + II + NO). We next considered different types of connections between graph A and either the R1 or R2 graph (C0-C8 graphs in Fig. 1). These connections range from no connection between allometric and reproductive traits (graph C0 in Fig. 1), to all three allometric traits affecting all six reproductive traits (graph C8 in Fig. 1).

To limit the number of possible trait networks and facilitate model interpretability, we did not consider direct connections between dietary and reproductive traits and we always included adult mass in all connection graphs (except C0). The response variable (Y; either population density or *N_e_*) was modeled as a function of all eleven predictor variables, equivalent to the M12 model in González-Suárez et al. (2021)^39^, or a subset of predictor variables equivalent to M3 and M8 models in González-Suárez et al. (2021)^39^ (Fig. 1). These three models were chosen because they represent the most frequently supported model structures reported by González-Suárez et al. (2021)^39^, finally resulting in 54 predictor-response networks considered (Supplementary Figs. 3-8). Both predictor and response variables were log_10_-transformed (except for dietary traits which were transformed by first taking the square root followed by the arcsine transformation) and standardized (transformed into z-scores) prior to the inference of regression coefficients in order to be able to compare coefficient estimates between models. When considering the “energetic equivalence” rule (EER)^33^ (Fig. 6), we only log_10_-transformed population density, adult mass and metabolic rate (without z-standardization) in order to be able to compare regression coefficients to previous studies.

To infer regression coefficients for the relationship between predictor and response variables, we used the “phylopath”^57^, “phylolm”^58^ and “daggity”^59^ libraries implemented in the R programming language. First, we tested whether the conditional independencies implied by each predictor–response network were supported by the data by calculating the corresponding Fisher’s C statistic^60^ and its associated *P*-value for each network, as implemented in the “phylo_path” function of the “phylopath” R package. A significant *P*-value of < 0.05 implies that the proposed causal structure is not supported by the observed variation in the data, and therefore all such networks were excluded from further consideration. Additionally, if the fit of the model produced warnings, such as convergence failures or non-identifiable parameters, the corresponding networks were also discarded. This procedure was conducted for 1,000 iterations of the mammalian phylogenetic tree to account for variance in phylogenetic relationship inference, as well as 3 different models of trait evolution (Pagel’s A, K and 0)^61^. In total, this amounted to 162,000 (54×1,000×3) evaluated predictor–response networks for each data subset (Supplementary Tables 2-21; see below for details on data subsetting).

We estimated regression coefficients representing the total effect of predictors on response variables for each combination of the predictor–response network, phylogenetic tree, and trait evolution model that passed the above-mentioned criteria. Additionally, if multiple models were supported for a specific data subset and phylogenetic tree, we compared their model fit using the C statistic Information Criterion (CIC_c_)^46^, similar to the classic Akaike Information Criterion^62^. The CIC_c_ allows for comparison of non-nested models^63^, with lower CIC_c_ values indicating better relative model support. We used the CIC_c_-based model rank to determine the best fitting model for every iteration of the phylogenetic tree, data subset and response variable. In the main text, we report regression coefficients estimated based on the most frequently supported model with respect to the assumed evolutionary trait model across all phylogenetic trees. Estimates of regression coefficients based on all supported models, along with their associated *P*-values, are reported in Supplementary Tables 22-49, while the distributions of supported predictor-response networks for each combination of data subset and response variable are reported in Supplementary Table 50. All analysis scripts are available at https://github.com/jbergman/mammal_traits.

### Data subsetting

Regression coefficients were estimated by considering all mammal species together, or separately based on three species classifications. First, we ran the coefficient estimation separately for species in three phylogenetic orders: Cetartiodactyla (ungulates; 99 species), Primates (primates; 83 species) and Carnivora (carnivorans; 60 species). Second, we ran coefficient estimation separately for species within three different trophic levels as defined in the COMBINE database: herbivore (191 species), omnivore (104 species) and carnivore (85 species). Third, we ran the coefficient estimation separately for species whose present-natural range^64^ (i.e., expected potential range without anthropogenic range restrictions) was predominantly within one of three biomes^65^, following the procedure in Bergman *et al*. (2023)^47^: cold (51 species), arid (106 species) and tropical (156 species).

## Results

### Trait Effects on Mammalian Population Density

We first considered the effects of phenotypic traits on mammalian population density estimated for current core areas of each species and thus assumed to be a proxy for natural densities^50^ (350 species with available density estimates). For every iteration of the mammalian phylogenetic tree, we fit 162 phylogenetic path models that differed with respect to the underlying predictor-response network (Fig. 1; 54 networks in total) and trait evolution model (3 models in total: Pagel’s A, K and 0-based model)^61^. For each model, we tested whether the conditional independencies between variables were upheld given each of the considered predictor-response networks and identified the best-fitting model for each tree (Supplementary Table 2; see Materials and Methods)^46^. Across most trees (98.7%), models assuming trait evolution under Pagel’s K transformation were ranked as the best-fitting. Therefore, to maintain consistency across analyses, we describe phylogenetic regression coefficients estimated using the K-based models in further text (Fig. 2). Regression coefficients estimated using the A-based models, which were broadly consistent with those obtained under the K-based model, are reported in Supplementary Fig. 9 and Supplementary Table 22. Across the 1,000 different iterations of the mammalian phylogenetic tree, supported K-based models were available for 993 trees and were distributed across two predictor-response networks: M12+C8+R2+A (95% of models; Supplementary Table 50) and M8+C8+R2+A (5% of models; Supplementary Table 50). The M12+C8+R2+A represents the most complex predictor-response network (Supplementary Fig. 8I). It incorporates the complete set of paths between reproductive variables (graph R2 in Fig. 1) by assuming that reproductive traits depend on all three allometric traits (adult mass, brain mass, and metabolic rate; graph C8 in Fig. 1), and includes a direct dependence between all traits and the response variable (graph M12 in Fig. 1). The M8+C8+R2+A network is only slightly less complex, with the difference that adult mass has no direct effect on population density, but rather acts through intermediary traits.

**Figure 2.**
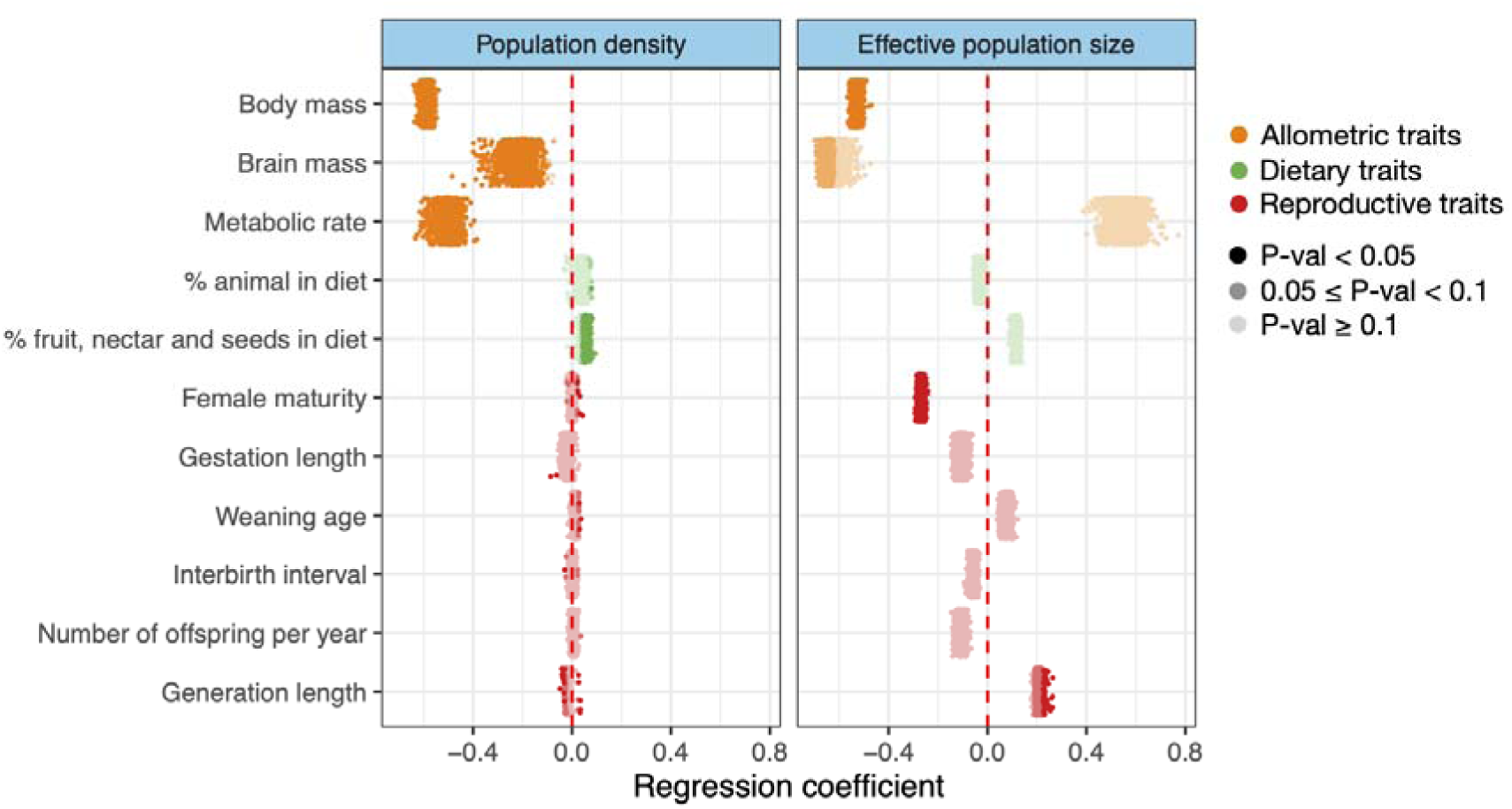
Estimates of regression coefficients for the total effects of phenotypic traits on population density (left panel; Supplementary Table 22) and effective population size (right panel; Supplementary Table 23) for terrestrial mammals and 1,00 iterations of the mammalian phylogenetic tree, given the most frequently supported model (Supplementary Table 50). Each point is a single regression coefficient corresponding to a single iteration of the phylogenetic tree, with color intensity representin statistical significance. The red dashed line at zero represents no effect of the predictor on the response variable.

Among allometric traits, adult mass had the strongest negative effect on density as observed previously^32–35^ and expected given the method of mass-dependent imputation used to infer this parameter^50^. The mean regression coefficient across all considered models was estimated to be −0.59 (range of 95% confidence intervals across all considered models: [−0.69, −0.49]). All estimated mass-related coefficients were statistically significant (*P*-value < 0.05). Metabolic rate had a significant negative effect on density across all models, but was slightly weaker compared to the mass effect (mean regression coefficient of −0.50 [−0.68, −0.29]). Brain mass had a negative effect on density that was largely significant (with *P*-value < 0.05 for 99% of models and and 0.05 ≤ *P*-value < 0.1 across 1% of models), but weaker still in magnitude compared to the other two allometric traits (mean regression coefficient of - 0.21 [−0.53, 0.05]).

Diet and reproduction had largely non-significant effects on density, with only the percentage of diet made up by fruit, nectar and seeds having a significantly positive impact (*P*-value < 0.05) on population density across 50% of considered models and a marginally significant effect (0.05 ≤ *P*-value < 0.1) across 33% of models (mean regression coefficient of 0.05 [−0.05, 0.14]). While the negative effect of allometry reflects the well-studied hypothesis that larger species have higher energy requirements that lead to lower population densities^33,36^, the marginally positive effects of a frugivorous diet may suggest that higher population densities are achieved in species with more complex foraging behaviors. Reproduction had a largely negligible effect on population density, with over 90% of models showing no significant effects of any reproductive trait on population density.

### Trait Effects on Mammalian *N_e_*

To investigate how phenotypic traits shape effective population sizes (*N_e_*), we calculated the long-term average of this parameter for 380 species of mammals. Similarly to the results obtained when using population density as the response variable, we found that -based models provided the best fit for th majority of phylogenetic trees (66%) when considering *N_e_* as the response variable (Supplementary Table 3). Results from the alternative -based models are presented in Supplementary Fig. 9 and Supplementary Table 23. The supported -based models (available for 998 trees) were distributed across three predictor–response networks: M12+C8+R2+A (1% of models; Supplementary Fig. 8I and Supplementary Table 50), M8+C8+R2+A (93% of models; Supplementary Fig. 6I and Supplementary Table 50) and M3+C8+R2+A (6% of models; Supplementary Fig. 4I and Supplementary Table 50). As with population density, a single predictor-response network was dominant: in the case of *N_e_*, the medium complexity graph for the relationship between traits and the response variable was most frequently included into th best-fitting models (graph M8 in Fig. 1).

As observed for population density, the effects of allometric traits had generally larger magnitude impacts compared to other traits (Fig. 2). However, only adult mass had a significantly negative effect on *N_e_* in all considered models (mean regression coefficient of −0.53 [−0.82, −0.20]), while brain mass had only a marginally negative effect (with 0.05 ≤ *P*-value < 0.1 across 47% of models; mean regression coefficient of −0.62 [−1.44, 0.28]), and a non-significant effect was observed for metabolic rate across all models. While we found no significant effect of dietary traits on *N_e_*, time to female maturity showed a significantly negative effect on *N_e_*across all models (mean regression coefficient of −0.27 [−0.49, −0.04]), indicating that species with shorter maturation periods tended to have higher effective sizes, even after allometric effects were taken into account. Additionally, the effect of generation length on *N_e_* wa positive, but mostly marginal (*P*-value < 0.05 across 6% of models and 0.05 ≤ *P*-value < 0.1 across 91% of models; mean regression coefficient of 0.21 [−0.05, 0.50]).

Taken together, these observations show that while population density and *N_e_* are largely affected by phenotypic traits in a similar manner, they can also be asymmetrically affected by the same phenotypic trait. As specific levels of both population density and *N_e_* need to be maintained in order to ensure species survival, the observed trait effects indicate the existence of evolutionary trade-offs between the two metrics of population dynamics.

### Order-specific patterns

Given that general patterns likely depend on species composition, we next focused on modelling population dynamics in three distinct mammalian orders with the largest number of species within our dataset: Cetartiodactyla (ungulates; 99 species), Primates (primates; 82 species) and Carnivora (carnivorans; 60 species). Model selection and regression coefficient inference was conducted separatel for each order. Models assuming trait evolution under Pagel’s transformation were most frequentl identified as the best-fitting (Supplementary Tables 4-9) and therefore presented here. Regression coefficients for all supported models are shown in Supplementary Figs. 10-13 and Tables 24-29, and the distribution of preferred predictor–response networks is provided in Supplementary Table 50.

Allometric traits had negative effects across all orders and response variables. We observed significantly negative associations (*P*-value < 0.05) between population density and both adult mass and metabolic rate, while brain size had a marginal effect. Ungulates displayed the strongest negative effect (*P*-value < 0.05 across 21% of models and 0.05 ≤ *P*-value < 0.1 across 16% of models; mean regression coefficient of −0.18 [−0.61, 0.24]). Conversely, the effect of allometry on *N_e_* was generally non-significant across orders, with notable exceptions. Ungulate *N_e_*was marginally negatively affected by adult mass (*P*-value < 0.05 across 16% of models and 0.05 ≤ *P*-value < 0.1 across 39% of models; mean regression coefficient of −0.26 [−0.63, 0.18]), while carnivoran *N_e_* was marginally positively affected by metabolic rate (*P*-value < 0.05 across 2% of models and 0.05 ≤ *P*-value < 0.1 across 22% of models; mean regression coefficient of 1.29 [−1.00, 3.81]). However, the strongest relationship between allometry and *N_e_*was evident in the positive effect of brain size on carnivoran *N_e_*(*P*-value < 0.05 across 92% of models and 0.05 ≤ *P*-value < 0.1 across 8% of models; mean regression coefficient of 1.43 [−0.23, 3.03]), indicating that the benefit of having a larger brain outweighs the higher energy requirements of brain tissue in this order (Fig. 3A).

**Figure 3.**
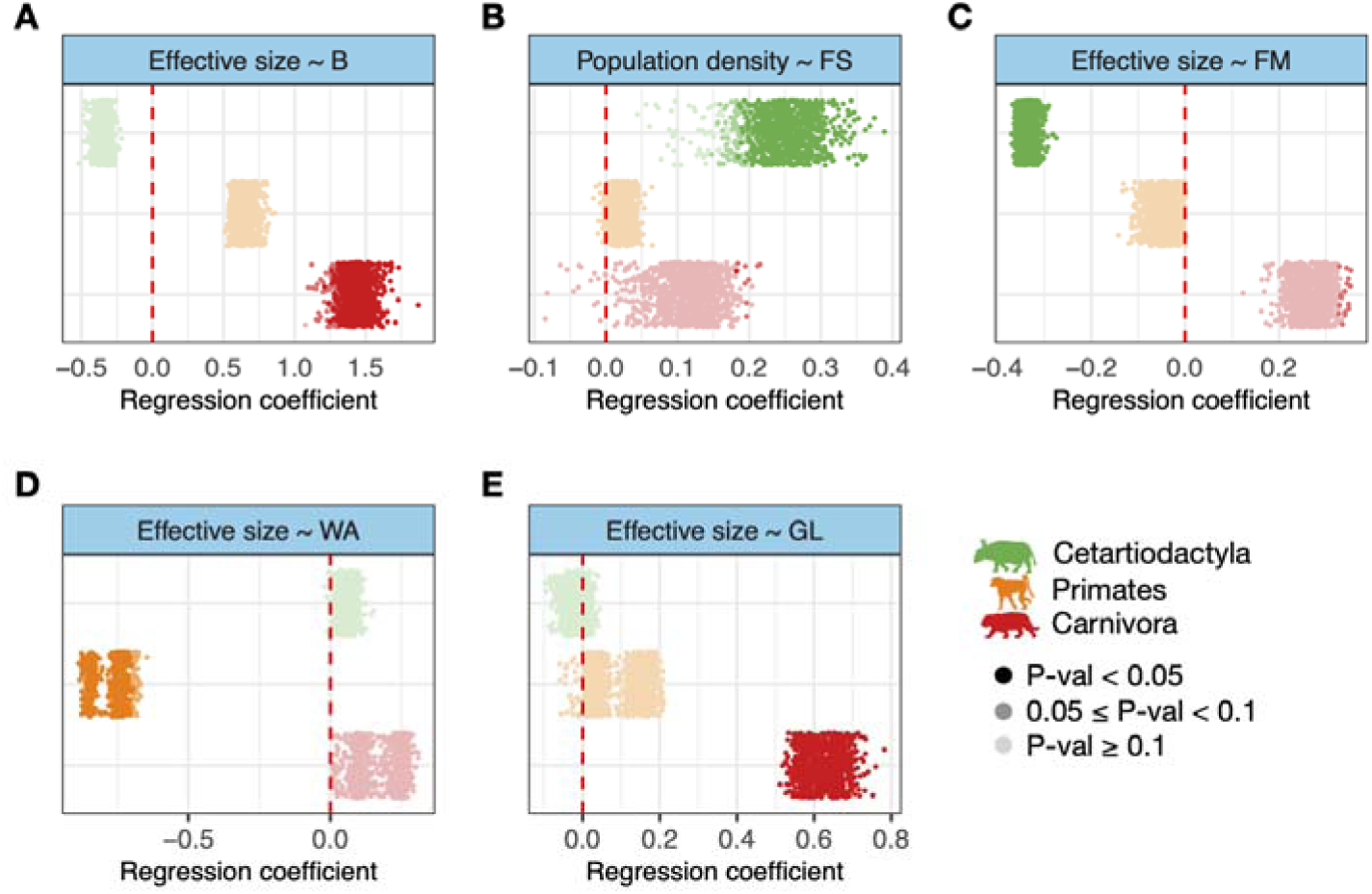
Order-specific regression coefficients for the total effects of **A.** brain mass (B), **B.** percentage of diet made up by fruit, nectar and seeds (FS), **C.** time to female maturity (FM), **D.** weaning age (WA) and **E.** generation length (GL) on effective population size (**A.**,**C.**-**E.**) and population density (**B.**) for terrestrial mammals. Coefficients were estimated separately for each order across 1,000 iterations of the mammalian phylogenetic tree. Color intensity indicates statistical significance. The re dashed line at zero represents no effect of the predictor on the response variable.

Dietary traits had variable effects across orders and response variables. In ungulates, the positive effect of a frugivorous diet on population density (*P*-value < 0.05 across 84% of models and 0.05 ≤ *P*-value < 0.1 across 11% of models; mean regression coefficient of 0.24 [−0.11, 0.58]) may indicate that a more diverse diet is a benefit to species in this order (Fig. 3B). Given that ungulates comprise the largest subset in our data and that the effects of dietary traits on primate and carnivoran population density are largely non-significant, the same positive effect observed in the general patterns (Fig. 2) is likely driven primarily by ungulates. Conversely, the effect of diet on *N_e_*was largely non-significant across all orders (Supplementary Figs. 10-13).

The effect of reproductive traits on population density was generally weak across orders, with a marginal effect on density in primates and carnivorans: a positive effect of female maturity in primates (*P*-value < 0.05 across 6% of models and 0.05 ≤ *P*-value < 0.1 across 72% of models; mean regression coefficient of 0.14 [−0.06, 0.31]), a negative effect of gestation length in primates (*P*-value < 0.05 across 4% of models 0.05 ≤ *P*-value < 0.1 across 63% of models; mean regression coefficient of −0.16 [−0.39, 0.11]), a positive effect of gestation length in carnivorans (*P*-value < 0.05 across 23% of models and 0.05 ≤ *P*-value < 0.1 across 28% of models; mean regression coefficient of 0.07 [−0.04, 0.19]) and a positive effect of weaning age in carnivorans (*P*-value < 0.05 across 25% of models and 0.05 ≤ *P*-value < 0.1 across 26% of models; mean regression coefficient of 0.07 [−0.06, 0.19]).

Conversely, the association between reproduction and *N_e_*yielded stronger patterns. First, time to female maturity in ungulates was significantly negatively associated with *N_e_* across all considered models (mean regression coefficient of −0.34 [−0.61, −0.03]), while no significant association was detected for the other two orders (Fig. 3C). The benefit of having a short maturation time may reflect the evolution of the precocial life history strategy in ungulates (production of a small number of highly developed offspring), hypothesized to have evolved as a means to escape predation^66^, which may in turn also shape ungulate *N_e_*.

Length of the the interbirth interval (*P*-value < 0.05 across 13% of models and 0.05 ≤ *P*-value < 0.1 across 61% of models; mean regression coefficient of −0.58 [−1.38, 0.06]), and particularly the weaning age in primates (*P*-value < 0.05 across 95% of models and 0.05 ≤ *P*-value < 0.1 across 5% of models; mean regression coefficient of −0.76 [−1.60, 0.06]), showed a negative association with *N_e_* (Fig. 3D). This observation indicates that longer weaning age in primates, and more generally, the extended juvenile period of primates, hypothesized to have evolved alongside increased behavioural complexity^67^, may limit effective population sizes in this order. While weaning age had no significant impact on ungulate or carnivoran *N_e_*, the length of the interbirth interval in ungulates showed the opposite trend to that of primates (*P*-value < 0.05 across 14% of models and 0.05 ≤ *P*-value < 0.1 across 76% of models; mean regression coefficient of 0.31 [−0.30, 0.77]), suggesting contrasting reproductive strategies between these groups.

Lastly, generation length had a positive impact on carnivoran *N_e_*across all considered models (mean regression coefficient of 0.63 [0.07, 1.17]), while no significant association was detected for the other two orders (Fig. 3E). The strong impact of generation length on carnivoran *N_e_* likely drives the corresponding trend observed in the general patterns (Fig. 2). This pattern may also suggest that longer generations promote carnivoran *N_e_*by allowing more time for dispersal and outbreeding in this order, or may reflect the evolution of altriciality (where offspring are born underdeveloped and require a longer time to reach maturity) in carnivorans.

### Patterns related to trophic levels and biome association

As demonstrated, subdivision of species yielded group-specific trait effects (Fig. 3) that were not observed at the global level (Fig. 2). In this section, we therefore considered other classifications of mammals, with respect to trophic levels and biome association to provide further insight into how life history context shapes species’ population dynamics. As before, we present regression coefficients estimated using the majority-preferred model (Supplementary Tables 10-21), while estimates obtained under all supported models are reported in Supplementary Figs. 14-25 and Supplementary Tables 30-41. The distributions of preferred predictor-response networks for combinations of species and response variables are presented in Supplementary Table 50.

We first investigated the effect of traits on population dynamics separately for species classified as herbivores, omnivores and carnivores. Here, we focused on the effect of dietary traits as they should b most clearly related to the classification of species with respect to their trophic level. We observed that the percentage of diet made up by animal items had a marginally positive effect on the population densit of herbivores (*P*-value < 0.05 across 2% of models and 0.05 ≤ *P*-value < 0.1 across 13% of models; mean regression coefficient of 0.05 [−0.03, 0.12]), whereas the opposite effect was observed for carnivores (*P*-value < 0.05 across 4% of models and 0.05 ≤ *P*-value < 0.1 across 15% of models; mean regression coefficient of −0.14 [−0.34, 0.08]), while no trend was observed for omnivores (Fig. 4A). A similar, but non-significant trend can be observed when considering *N_e_* as the response variable (Supplementary Figs. 14-19). Similarly, the percentage of diet made up by fruit, nectar and seeds had a marginally positive effect on herbivore population density (*P*-value < 0.05 across 18% of models and 0.05 ≤ *P*-value < 0.1 across 28% of models; mean regression coefficient of 0.06 [−0.09, 0.23]), while no effect was observed for the other two classifications (Fig. 4B). Taken together, these results suggest that dietary specialization has a modest negative effect on population dynamics, with dietary diversity having the most notable effect on herbivore density.

**Figure 4.**
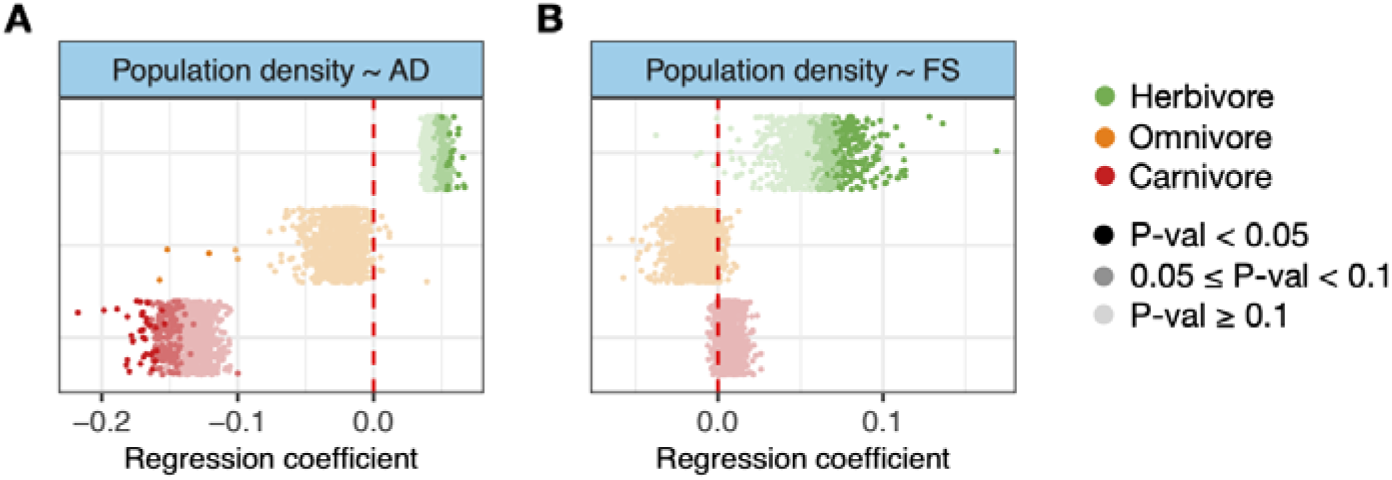
Trophic level-specific estimates of regression coefficients for the total effect of **A.** percentage of diet made up by animal items (AD) and **B.** percentage of diet made up by fruit, nectar and seeds (FS) on population density for terrestrial mammals. Coefficients were estimated separately for each order across 1,000 iterations of the mammalian phylogenetic tree. Color intensity indicates statistical significance. The red dashed line at zero represents no effect of the predictor on the response variable.

When subdividing species in our dataset by biome, we found several patterns that may reflect adaptive strategies of species to different habitats. First, while adult mass showed a significant, negative association with both population density and *N_e_* across all biomes and models considered (Supplementary Figs. 20-25), this association was strongest for species occupying cold biomes. A trend of association strength across biomes is especially evident in the relationship between adult mass and *N_e_* (Fig. 5A), with cold-adapted species showing the strongest association (mean regression coefficient of −1.04 [−1.56, - 0.51]), followed by species occupying arid biomes (mean regression coefficient of −0.76 [−1.21, −0.28]) and finally species within tropical biomes (mean regression coefficient of −0.31 [−0.55, −0.09]). This observation indicates that the evolution of larger body size as a means of adaptation to cold climates^68^ has had a disproportionately stronger negative effect on the *N_e_* of cold-adapted species.

**Figure 5.**
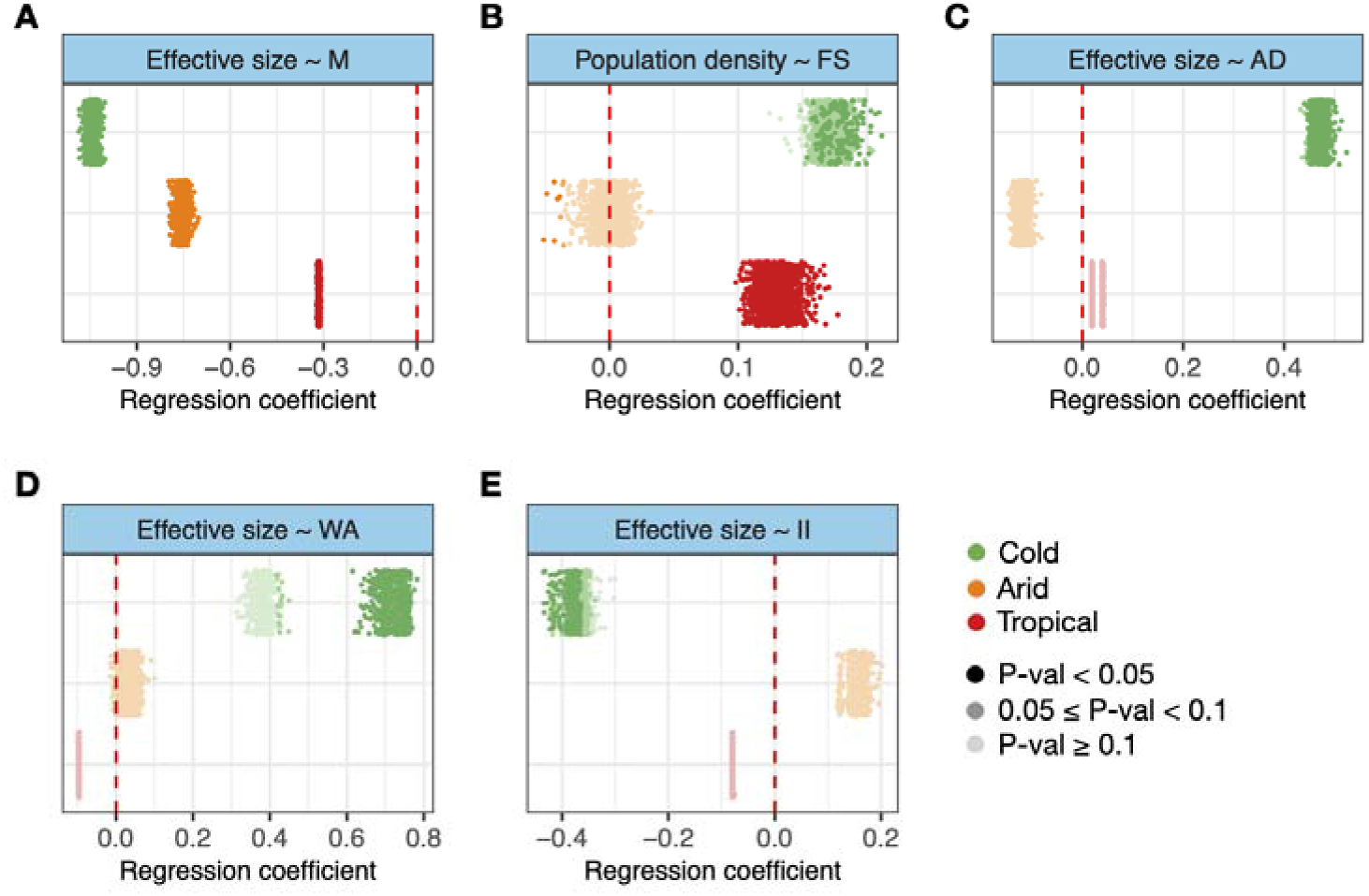
Biome-specific regression coefficients for the total effect of **A.** adult mass (M) **B.** percentage of diet made up by fruit, nectar and seeds (FS), **C.** percentage of diet made up by animal items (AD), **D.** weaning age (WA) and **E.** length of interbirth interval (II) on effective population size (**A.**,**C.**-**E.**) and population density (**B.**) for terrestrial mammals. Coefficients were estimated separately for each order across 1,000 iterations of the mammalian phylogenetic tree. Color intensity indicates statistical significance. The red dashed line at zero represents no effect of the predictor on the response variable.

We also observed notable effects of dietary traits when subdividing species with respect to th biome occupied. Population density of cold-adapted species showed a marginal, positive association with a frugivorous diet (*P*-value < 0.05 across 16% of models and 0.05 ≤ *P*-value < 0.1 across 77% of models; mean regression coefficient of 0.17 [−0.08, 0.42]; Fig. 5B). In contrast, the *N_e_* of cold-adapted species was positively associated with a carnivorous diet (*P*-value < 0.05 across 80% and 0.05 ≤ *P*-value < 0.1 acros 20% of models; mean regression coefficient of 0.47 [−0.04, 0.99]; Fig. 5C). For species occupying th tropical biome, a frugivorous diet there was significantly and positively associated with both population density (mean regression coefficient of 0.13 [0.02, 0.26]; Fig. 5B) and *N_e_* (mean regression coefficient of 0.22 [0.02, 0.43]; Supplementary Fig. 24) across all models. These associations may reflect differences in biome-specific resource availability, but also dietary adaptations of species that occupy each biome.

Several reproductive traits had notable effects on the population dynamics of cold-adapted species. Firstly, the population density of species occupying cold biomes was negatively but marginally associated with generation length (*P*-value < 0.05 across 23% of models and 0.05 ≤ *P*-value < 0.1 acros 64% of models; mean regression coefficient of −0.14 [−0.38, 0.04]; Supplementary Fig. 20). The *N_e_* of cold-adapted species was positively associated with weaning age (*P*-value < 0.05 across 69% of model and 0.05 ≤ *P*-value < 0.1 across 2% of models; mean regression coefficient of −0.14 [−0.38, 0.04]; Fig, 5D), but negatively associated with the length of the interbirth interval (*P*-value < 0.05 across 46% of models and 0.05 ≤ *P*-value < 0.1 across 31% of models; mean regression coefficient of −0.37 [−0.88, 0.19]; Fig. 5E). As these associations are not present for species occupying any of the other two biomes, they may reflect a unique adaptive strategy to colder climates, characterized by longer postnatal care coupled with relatively shorter intervals between reproductive events. We also found unique and significant associations between time to female maturity and the *N_e_* of species occupying the arid biome (mean regression coefficient of −0.53 [−0.89, −0.13]; Supplementary Fig. 22), and between gestation time and the *N_e_* of tropical species (mean regression coefficient of 0.18 [0.003, 0.36]; Supplementary Fig. 24), which may reflect biome-specific features or the ecological composition of species occupying these biomes.

### Population density and energetic equivalence

One of the most influential investigations of trait impacts on population dynamics is the study of the relationship between species’ population density and adult mass by Damuth (1981)^33^. Damuth showed that larger species achieve lower densities with the slope of the density-mass relationship estimated to be - 0.75, while the slope of the effect of mass on metabolic rate was exactly reciprocal (0.75; Kleiber’ law^69^). This led to the proposition of the “energetic equivalence” rule (EER)^33,36^, implying that energy requirements of a population are independent of adult mass. We tested this proposition for the species in our dataset by comparing the regression coefficient for the effect of adult mass on population density to the regression coefficient estimated for the effect of adult mass on metabolic rate. We found that adult mass had a smaller absolute effect on density compared to its effect on metabolic rate (Fig. 6), thus not supporting the EER hypothesis and showing that larger species use proportionally more resources within their environments. Specifically, we found that the expected reciprocity between density-mass and metabolic rate-mass relationships, assessed as the overlap in the absolute 95% confidence intervals of their regression coefficients, was not supported across any of the considered models or order-specific species subdivisions (Supplementary Tables 42-49). This is in line with previous studies that similarly took phylogenetic relatedness into account when assessing density-allometry relationships^35,39^.

**Figure 6.**
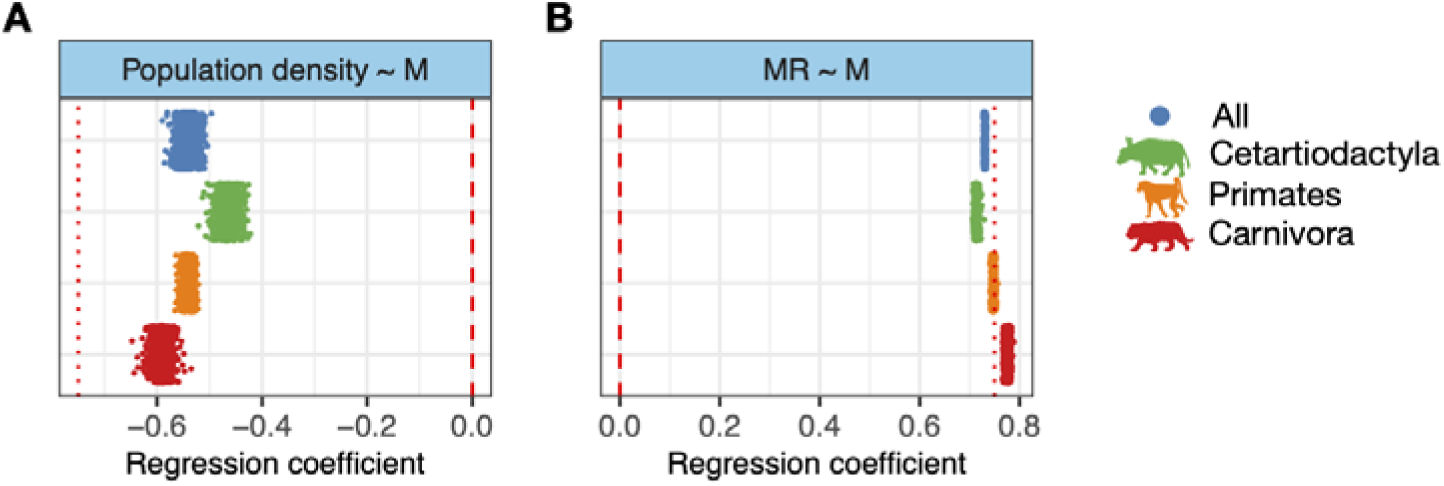
Estimates of regression coefficients for the total effect of adult mass (M) on **A.** population density and **B.** metabolic rate (MR) when considering either all mammal species together or different orders of terrestrial mammals, and 1,000 iterations of the mammalian phylogenetic tree. Each point is a single regression coefficient corresponding to a single iteration of the phylogenetic tree. All estimates of regression coefficients are significant at *P*-value < 0.05. The red dashed line at zero represents no effect of the predictor on the response variable, while dotted lines represent regression coefficients of 0.75 and −0.75 predicted by the “energetic equivalence” rule.

## Discussion

In this work, we show that mammalian life history strategies uniquely shape the relationship between traits and key population characteristics. By explicitly contrasting trait dependence of population density and long-term effective population size, our study identifies structured divergences between ecological and evolutionary population dynamics that are not apparent when either metric is considered in isolation.

We corroborated the widely observed negative impact of adult mass on population density^32–35^, while the association between mass and long-term effective population size was weak or context dependent (Fig. 2, Supplementary Figs. 9-25). This decoupling mirrors macroevolutionary evidence that large body size does not necessarily entail higher long-term extinction risk, but can instead be associated with lineage persistence^70^ through higher-level processes such as spatial connectivity, demographic buffering, or differential survival. Together, these findings suggest that the long-term evolutionary consequences of body size are mediated by processes operating beyond local ecological constraints. Furthermore, we found that effective population size, but not density, was related to brain mass, and likely to complexity of behavioral patterns, especially in carnivorans (Fig. 3)^71,72^. Conversely, larger brain size did not seem to be advantageous in other orders, which in the case of ungulates could be due to relatively restricted drivers of brain size evolution in these species^73^. The positive effect of brain size on *N_e_* further implies that species with larger brains occupy relatively large geographic ranges (resulting in lower density), in line with large-brained species being more adaptable to different environments, but are still able to maintain reproductive connectivity.

We also found that dietary specialization was negatively associated with population dynamics, especially affecting densities of herbivorous mammals (Fig. 3B and 4). This may reflect lower resource availability for species that are either highly specialized in their diet or occupy higher trophic levels within their home range^74,75^. On the other hand, the negative association of population dynamics and diet specialization, together with the relatively weak density-body mass scaling in ungulates (Fig. 6), suggests that large generalist mammals may achieve relatively higher densities than expected for their size, making them promising candidates for restoration efforts^76,77^.

*N_e_* was associated with reproductive strategies of mammals - specifically, ungulate precociality (Fig. 3C) and carnivoran altriciality (Fig. 3E) - implying that while they differ in intrinsic rates of growth^78^, both reproduction strategies have similar impacts on *N_e_*. In primates, the relationship between population dynamics and reproductive traits reflected their extended juvenile period^67^, likely a key adaptation in this order (Fig. 3D). We also observed a gradient of biome-specific relationships between *N_e_* and adult mass, as well as other distinct characteristics of cold-adapted species (Fig. 5), likely reflective of their unique evolutionary adaptations and life history^79–81^.

Different aspects of population dynamics underlie specific mechanisms that can ultimately result in species extinction. For example, low population densities can increase the vulnerability of a population to predation^82^ and/or decrease the probability of mate finding^83,84^. Similarly, low *N_e_* values can be a sign of inbreeding depression in a population^85^ and/or a decreased potential to adapt to future environmental change^86^. Thus, both population density and effective population size are required to be maintained at sufficient levels to ensure the successful propagation of a species. However, the relative importance of achieving a certain density or a certain *N_e_* value will likely vary between species and populations. Thus, species survival is likely to depend on the limiting factor of population dynamics, and may ultimately result in evolutionary trade-offs between different aspects of population dynamics. Our analyses demonstrate that such trade-offs are indeed observable, implying that evolution of life history strategies resulted in trade-offs not only at the trait level, but also at higher levels of biological organization, such as population characteristics. This observation challenges the static nature of the “energetic equivalence” rule (Fig. 6) proposed by initial studies of the density-allometry relationship^33,36^. Together with numerous previous criticisms^34,35,37,87–90^, we warn against overly simplistic interpretations of allometric scaling in ecology and emphasize the need for a more nuanced understanding of how these relationships vary across different species and contexts. Additionally, recent work exemplifies the use of non-linear relationships when considering life history traits^91^. It is also important to note that trait evolution may have occurred during the time period over which *N_e_*was estimated, which represents a potential limitation of our approach. Future work could address this limitation by using fossil-informed or otherwise time-calibrated trait reconstructions, to better align trait values with metrics of population dynamics.

In addition to estimates of population density, population genetic parameters are becoming increasingly available for a variety of species^92^, and are thus predicted to become widely used metrics for assessing population health and guiding species conservation and restoration^93^. However, it is important to note that SMC-based methods estimate the instantaneous coalescent rate^94^, the inverse of which corresponds to *N_e_*in panmictic populations - consequently, unaccounted population structure, as well as low genomic read coverage and uncertainty in mutation rate estimates (required to express SMC output in units of time and effective population size), both in terms of the approach used to estimate the rate^95^ and the possibility of rate change over evolutionary timescales^96,97^, can bias inferred trajectories^94,98–100^. Given that these sources of bias are typically most pronounced at a single-species level, their overall impact is likely reduced in our comparative framework, which spans hundreds of mammalian species for which high-quality genomic data are available. The average genomic read coverage across species in our dataset is 59× (Supplementary Table 1), a level that substantially reduces the likelihood of technical biases associated with low data quality^98^. We also estimated *N_e_* for each species as an average across a broad time window (100–800 kya), rather than focusing on short-term fluctuations across the full SMC-inferred trajectory. This approach provides a more robust and comparable summary of long-term effective population size, minimizing the influence of short-term estimation noise and avoiding biases at recent epochs that may arise from limitations of SMC methods and/or potential demographic impacts associated with humans^47,101,102^. The chosen estimate of *N_e_* thus provides a comparable timeframe to life history evolution, as well as an informative proxy for estimating baseline census population sizes, biomass and energy turnover of pre-anthropogenic ecosystems^47,54^. Generally, population genetic parameters, such as long-term *N_e_*, are a better representation of the dynamics and conditions under which extant species evolved, which extend far beyond the time scale of anthropogenic landscapes. Importantly, while population density may be maintained at local levels, effective population size is likely too low for a large proportion of mammals^47,54^, which can be alleviated through strategic dispersal and breeding of individuals at the metapopulation level.

Understanding the relationship between life history and population dynamics is crucial to predict and mitigate biodiversity decline, given that species with different life history strategies are expected to react differently to future environmental shifts^103^. It is thus also expected that conservation and restoration strategies will differ across species. For example, while an increase of population densities may be prioritized for some species, maintaining effective population size may be more important in other cases, making it crucial to understand trade-offs between different aspects of population dynamics. Our study thus provides a starting point for developing better-informed conservation and restoration strategies that will ultimately ensure the maintenance and protection of our planet’s biodiversity.

## Acknowledgments

This work was supported by the Center for Ecological Dynamics in a Novel Biosphere (ECONOVO), funded by the Danish National Research Foundation (grant DNRF173 to J.-C.S.). All of the computing for this project was performed on the GenomeDK cluster (https://genome.au.dk/). We would like to thank GenomeDK and Aarhus University for providing computational resources and support that contributed to these research results.

## Supplement

### Supplementary Figure Legends

Supplementary Figure 1. Correlation plots between allometric, dietary and reproductive traits. All variables are log_10_-transformed and z-score standardized. Variables include adult mass (M), brain mass (B), metabolic rate (MR), percentage of diet made up by animal items (AD), percentage of diet made up by fruit, nectar and seeds (FS), time to female maturity (FM), gestation length (GS), weaning age (WA), length of interbirth interval (II), number of offspring per year (NO) and generation length (GL).

Supplementary Figure 2. Correlation plot between response variables, population density and effective population size (harmonic mean 100-800 kya). Both variables are log_10_-transformed and z-score standardized.

Supplementary Figure 3. The schematic representations of full predictor-response networks constructed from R1, M3 and **A.** C0, **B.** C1, **C.** C2, **D.** C3, **E.** C4, **F.** C5, **G.** C6, **H.** C7, **I.** C8 graphs.

Supplementary Figure 4. The schematic representations of full predictor-response networks constructed from R2, M3 and **A.** C0, **B.** C1, **C.** C2, **D.** C3, **E.** C4, **F.** C5, **G.** C6, **H.** C7, **I.** C8 graphs.

Supplementary Figure 5. The schematic representations of full predictor-response networks constructed from R1, M8 and **A.** C0, **B.** C1, **C.** C2, **D.** C3, **E.** C4, **F.** C5, **G.** C6, **H.** C7, **I.** C8 graphs.

Supplementary Figure 6. The schematic representations of full predictor-response networks constructed from R2, M8 and **A.** C0, **B.** C1, **C.** C2, **D.** C3, **E.** C4, **F.** C5, **G.** C6, **H.** C7, **I.** C8 graphs.

Supplementary Figure 7. The schematic representations of full predictor-response networks constructed from R1, M12 and **A.** C0, **B.** C1, **C.** C2, **D.** C3, **E.** C4, **F.** C5, **G.** C6, **H.** C7, **I.** C8 graphs.

Supplementary Figure 8. The schematic representations of full predictor-response networks constructed from R2, M12 and **A.** C0, **B.** C1, **C.** C2, **D.** C3, **E.** C4, **F.** C5, **G.** C6, **H.** C7, **I.** C8 graphs.

Supplementary Figure 9. Estimates of regression coefficients for the total effects of phenotypic traits on population density (left panel; Supplementary Table 22) and effective population size (right panel; Supplementary Table 23) for all terrestrial mammals and 1,000 iterations of the mammalian phylogenetic tree, given the alternatively supported model (Supplementary Table 50). Each point is a single regression coefficient corresponding to a single iteration of the phylogenetic tree, with increasing degrees of color intensity representing statistical significance. The red dashed line at zero represents no effect of the predictor on the response variable.

Supplementary Figure 10. Estimates of regression coefficients for the total effects of phenotypic traits on population density (left panel; Supplementary Table 24) and effective population size (right panel; Supplementary Table 25) restricted to Cetartiodactyla species and 1,000 iterations of the mammalian phylogenetic tree, given the most frequently supported model (Supplementary Table 50). Each point is a single regression coefficient corresponding to a single iteration of the phylogenetic tree, with increasing degrees of color intensity representing statistical significance. The red dashed line at zero represents no effect of the predictor on the response variable. Note that estimates for Cetartiodactyla species given the alternatively supported model are not available as only a single model type was supported (Supplementary Table 50).

Supplementary Figure 11. Estimates of regression coefficients for the total effects of phenotypic traits on population density (left panel; Supplementary Table 26) and effective population size (right panel; Supplementary Table 27) restricted to Primates species and 1,000 iterations of the mammalian phylogenetic tree, given the most frequently supported model (Supplementary Table 50). Each point is a single regression coefficient corresponding to a single iteration of the phylogenetic tree, with increasing degrees of color intensity representing statistical significance. The red dashed line at zero represents no effect of the predictor on the response variable.

Supplementary Figure 12. Estimates of regression coefficients for the total effects of phenotypic traits on population density (left panel; Supplementary Table 26) and effective population size (right panel; Supplementary Table 27) restricted to Primates species and 1,000 iterations of the mammalian phylogenetic tree, given the alternatively supported model (Supplementary Table 50). Each point is a single regression coefficient corresponding to a single iteration of the phylogenetic tree, with increasing degrees of color intensity representing statistical significance. The red dashed line at zero represents no effect of the predictor on the response variable.

Supplementary Figure 13. Estimates of regression coefficients for the total effects of phenotypic traits on population density (left panel; Supplementary Table 28) and effective population size (right panel; Supplementary Table 29) restricted to Carnivora species and 1,000 iterations of the mammalian phylogenetic tree, given the most frequently supported model (Supplementary Table 50). Each point is a single regression coefficient corresponding to a single iteration of the phylogenetic tree, with increasing degrees of color intensity representing statistical significance. The red dashed line at zero represents no effect of the predictor on the response variable. Note that estimates for Carnivora species given the alternatively supported model are not available as only a single model type was supported (Supplementary Table 50).

Supplementary Figure 14. Estimates of regression coefficients for the total effects of phenotypic traits on population density (left panel; Supplementary Table 30) and effective population size (right panel; Supplementary Table 31) restricted to herbivore species and 1,000 iterations of the mammalian phylogenetic tree, given the most frequently supported model (Supplementary Table 50). Each point is a single regression coefficient corresponding to a single iteration of the phylogenetic tree, with increasing degrees of color intensity representing statistical significance. The red dashed line at zero represents no effect of the predictor on the response variable.

Supplementary Figure 15. Estimates of regression coefficients for the total effects of phenotypic traits on population density (left panel; Supplementary Table 30) and effective population size (right panel; Supplementary Table 31) restricted to herbivore species and 1,000 iterations of the mammalian phylogenetic tree, given the alternatively supported model (Supplementary Table 50). Each point is a single regression coefficient corresponding to a single iteration of the phylogenetic tree, with increasing degrees of color intensity representing statistical significance. The red dashed line at zero represents no effect of the predictor on the response variable.

Supplementary Figure 16. Estimates of regression coefficients for the total effects of phenotypic traits on population density (left panel; Supplementary Table 32) and effective population size (right panel; Supplementary Table 33) restricted to omnivore species and 1,000 iterations of the mammalian phylogenetic tree, given the most frequently supported model (Supplementary Table 50). Each point is a single regression coefficient corresponding to a single iteration of the phylogenetic tree, with increasing degrees of color intensity representing statistical significance. The red dashed line at zero represents no effect of the predictor on the response variable.

Supplementary Figure 17. Estimates of regression coefficients for the total effects of phenotypic traits on population density (left panel; Supplementary Table 32) and effective population size (right panel; Supplementary Table 33) restricted to omnivore species and 1,000 iterations of the mammalian phylogenetic tree, given the alternatively supported model (Supplementary Table 50). Each point is a single regression coefficient corresponding to a single iteration of the phylogenetic tree, with increasing degrees of color intensity representing statistical significance. The red dashed line at zero represents no effect of the predictor on the response variable.

Supplementary Figure 18. Estimates of regression coefficients for the total effects of phenotypic traits on population density (left panel; Supplementary Table 34) and effective population size (right panel; Supplementary Table 35) restricted to carnivore species and 1,000 iterations of the mammalian phylogenetic tree, given the most frequently supported model (Supplementary Table 50). Each point is a single regression coefficient corresponding to a single iteration of the phylogenetic tree, with increasing degrees of color intensity representing statistical significance. The red dashed line at zero represents no effect of the predictor on the response variable.

Supplementary Figure 19. Estimates of regression coefficients for the total effects of phenotypic traits on population density (left panel; Supplementary Table 34) and effective population size (right panel; Supplementary Table 35) restricted to carnivore species and 1,000 iterations of the mammalian phylogenetic tree, given the alternatively supported model (Supplementary Table 50). Each point is a single regression coefficient corresponding to a single iteration of the phylogenetic tree, with increasing degrees of color intensity representing statistical significance. The red dashed line at zero represents no effect of the predictor on the response variable.

Supplementary Figure 20. Estimates of regression coefficients for the total effects of phenotypic traits on population density (left panel; Supplementary Table 36) and effective population size (right panel; Supplementary Table 37) restricted to species occupying the cold biome and 1,000 iterations of the mammalian phylogenetic tree, given the most frequently supported model (Supplementary Table 50). Each point is a single regression coefficient corresponding to a single iteration of the phylogenetic tree, with increasing degrees of color intensity representing statistical significance. The red dashed line at zero represents no effect of the predictor on the response variable.

Supplementary Figure 21. Estimates of regression coefficients for the total effects of phenotypic traits on population density (Supplementary Table 36) restricted to species occupying the cold biome and 1,000 iterations of the mammalian phylogenetic tree, given the alternatively supported model (Supplementary Table 50). Each point is a single regression coefficient corresponding to a single iteration of the phylogenetic tree, with increasing degrees of color intensity representing statistical significance. The red dashed line at zero represents no effect of the predictor on the response variable. Note that estimates for effective population size as response given the alternatively supported model are not available as only a single model type was supported (Supplementary Table 50).

Supplementary Figure 22. Estimates of regression coefficients for the total effects of phenotypic traits on population density (left panel; Supplementary Table 38) and effective population size (right panel; Supplementary Table 39) restricted to species occupying the arid biome and 1,000 iterations of the mammalian phylogenetic tree, given the most frequently supported model (Supplementary Table 50). Each point is a single regression coefficient corresponding to a single iteration of the phylogenetic tree, with increasing degrees of color intensity representing statistical significance. The red dashed line at zero represents no effect of the predictor on the response variable.

Supplementary Figure 23. Estimates of regression coefficients for the total effects of phenotypic traits on population density (left panel; Supplementary Table 40) and effective population size (right panel; Supplementary Table 41) restricted to species occupying the arid biome and 1,000 iterations of the mammalian phylogenetic tree, given the alternatively supported model (Supplementary Table 50). Each point is a single regression coefficient corresponding to a single iteration of the phylogenetic tree, with increasing degrees of color intensity representing statistical significance. The red dashed line at zero represents no effect of the predictor on the response variable.

Supplementary Figure 24. Estimates of regression coefficients for the total effects of phenotypic traits on population density (left panel; Supplementary Table 40) and effective population size (right panel; Supplementary Table 41) restricted to species occupying the tropical biome and 1,000 iterations of the mammalian phylogenetic tree, given the most frequently supported model (Supplementary Table 50). Each point is a single regression coefficient corresponding to a single iteration of the phylogenetic tree, with increasing degrees of color intensity representing statistical significance. The red dashed line at zero represents no effect of the predictor on the response variable.

Supplementary Figure 25. Estimates of regression coefficients for the total effects of phenotypic traits on population density (left panel; Supplementary Table 38) and effective population size (right panel; Supplementary Table 39) restricted to species occupying the tropical biome and 1,000 iterations of the mammalian phylogenetic tree, given the alternatively supported model (Supplementary Table 50). Each point is a single regression coefficient corresponding to a single iteration of the phylogenetic tree, with increasing degrees of color intensity representing statistical significance. The red dashed line at zero represents no effect of the predictor on the response variable.

### Supplementary Table Legends

Supplementary Table 1. List of species used in the study including classifications, trait values and short read and reference genome accessions used in the bioinformatic pipeline.

Supplementary Table 2. Phylogenetic path model summaries for analyses including all species, with population density as the response variable, across all combinations of predictor-response networks, trait evolution models and phylogenetic trees. Model statistics include the the number of independence claims made by the model (k), the number of model parameters (q), Fisher’s C goodness-of-fit statistic (C) and associated P-value (p), and the small-sample-corrected C information criterion (CICc).

Supplementary Table 3. Phylogenetic path model summaries for analyses including all species, with effective population size as the response variable, across all combinations of predictor-response networks, trait evolution models and phylogenetic trees. Model statistics include the the number of independence claims made by the model (k), the number of model parameters (q), Fisher’s C goodness-of-fit statistic (C) and associated P-value (p), and the small-sample-corrected C information criterion (CICc).

Supplementary Table 4. Phylogenetic path model summaries for analyses restricted to Cetartiodactyla species, with population density as the response variable, across all combinations of predictor-response networks, trait evolution models and phylogenetic trees. Model statistics include the the number of independence claims made by the model (k), the number of model parameters (q), Fisher’s C goodness-of-fit statistic (C) and associated P-value (p), and the small-sample-corrected C information criterion (CICc).

Supplementary Table 5. Phylogenetic path model summaries for analyses restricted to Cetartiodactyla species, with effective population size as the response variable, across all combinations of predictor-response networks, trait evolution models and phylogenetic trees. Model statistics include the the number of independence claims made by the model (k), the number of model parameters (q), Fisher’s C goodness-of-fit statistic (C) and associated P-value (p), and the small-sample-corrected C information criterion (CICc).

Supplementary Table 6. Phylogenetic path model summaries for analyses restricted to Primates species, with population density as the response variable, across all combinations of predictor-response networks, trait evolution models and phylogenetic trees. Model statistics include the the number of independence claims made by the model (k), the number of model parameters (q), Fisher’s C goodness-of-fit statistic (C) and associated P-value (p), and the small-sample-corrected C information criterion (CICc).

Supplementary Table 7. Phylogenetic path model summaries for analyses restricted to Primates species, with effective population size as the response variable, across all combinations of predictor-response networks, trait evolution models and phylogenetic trees. Model statistics include the the number of independence claims made by the model (k), the number of model parameters (q), Fisher’s C goodness-of-fit statistic (C) and associated P-value (p), and the small-sample-corrected C information criterion (CICc).

Supplementary Table 8. Phylogenetic path model summaries for analyses restricted to Carnivora species, with population density as the response variable, across all combinations of predictor-response networks, trait evolution models and phylogenetic trees. Model statistics include the the number of independence claims made by the model (k), the number of model parameters (q), Fisher’s C goodness-of-fit statistic (C) and associated P-value (p), and the small-sample-corrected C information criterion (CICc).

Supplementary Table 9. Phylogenetic path model summaries for analyses restricted to Carnivora species, with effective population size as the response variable, across all combinations of predictor-response networks, trait evolution models and phylogenetic trees. Model statistics include the the number of independence claims made by the model (k), the number of model parameters (q), Fisher’s C goodness-of-fit statistic (C) and associated P-value (p), and the small-sample-corrected C information criterion (CICc).

Supplementary Table 10. Phylogenetic path model summaries for analyses restricted to herbivore species, with population density as the response variable, across all combinations of predictor-response networks, trait evolution models and phylogenetic trees. Model statistics include the the number of independence claims made by the model (k), the number of model parameters (q), Fisher’s C goodness-of-fit statistic (C) and associated P-value (p), and the small-sample-corrected C information criterion (CICc).

Supplementary Table 11. Phylogenetic path model summaries for analyses restricted to herbivore species, with effective population size as the response variable, across all combinations of predictor-response networks, trait evolution models and phylogenetic trees. Model statistics include the the number of independence claims made by the model (k), the number of model parameters (q), Fisher’s C goodness-of-fit statistic (C) and associated P-value (p), and the small-sample-corrected C information criterion (CICc).

Supplementary Table 12. Phylogenetic path model summaries for analyses restricted to omnivore species, with population density as the response variable, across all combinations of predictor-response networks, trait evolution models and phylogenetic trees. Model statistics include the the number of independence claims made by the model (k), the number of model parameters (q), Fisher’s C goodness-of-fit statistic (C) and associated P-value (p), and the small-sample-corrected C information criterion (CICc).

Supplementary Table 13. Phylogenetic path model summaries for analyses restricted to omnivore species, with effective population size as the response variable, across all combinations of predictor-response networks, trait evolution models and phylogenetic trees. Model statistics include the the number of independence claims made by the model (k), the number of model parameters (q), Fisher’s C goodness-of-fit statistic (C) and associated P-value (p), and the small-sample-corrected C information criterion (CICc).

Supplementary Table 14. Phylogenetic path model summaries for analyses restricted to carnivore species, with population density as the response variable, across all combinations of predictor-response networks, trait evolution models and phylogenetic trees. Model statistics include the the number of independence claims made by the model (k), the number of model parameters (q), Fisher’s C goodness-of-fit statistic (C) and associated P-value (p), and the small-sample-corrected C information criterion (CICc).

Supplementary Table 15. Phylogenetic path model summaries for analyses restricted to carnivore species, with effective population size as the response variable, across all combinations of predictor-response networks, trait evolution models and phylogenetic trees. Model statistics include the the number of independence claims made by the model (k), the number of model parameters (q), Fisher’s C goodness-of-fit statistic (C) and associated P-value (p), and the small-sample-corrected C information criterion (CICc).

Supplementary Table 16. Phylogenetic path model summaries for analyses restricted to species occupying the cold biome, with population density as the response variable, across all combinations of predictor-response networks, trait evolution models and phylogenetic trees. Model statistics include the the number of independence claims made by the model (k), the number of model parameters (q), Fisher’s C goodness-of-fit statistic (C) and associated P-value (p), and the small-sample-corrected C information criterion (CICc).

Supplementary Table 17. Phylogenetic path model summaries for analyses restricted to species occupying the cold biome, with effective population size as the response variable, across all combinations of predictor-response networks, trait evolution models and phylogenetic trees. Model statistics include the the number of independence claims made by the model (k), the number of model parameters (q), Fisher’s C goodness-of-fit statistic (C) and associated P-value (p), and the small-sample-corrected C information criterion (CICc).

Supplementary Table 18. Phylogenetic path model summaries for analyses restricted to species occupying the arid biome, with population density as the response variable, across all combinations of predictor-response networks, trait evolution models and phylogenetic trees. Model statistics include the the number of independence claims made by the model (k), the number of model parameters (q), Fisher’s C goodness-of-fit statistic (C) and associated P-value (p), and the small-sample-corrected C information criterion (CICc).

Supplementary Table 19. Phylogenetic path model summaries for analyses restricted to species occupying the arid biome, with effective population size as the response variable, across all combinations of predictor-response networks, trait evolution models and phylogenetic trees. Model statistics include the the number of independence claims made by the model (k), the number of model parameters (q), Fisher’s C goodness-of-fit statistic (C) and associated P-value (p), and the small-sample-corrected C information criterion (CICc).

Supplementary Table 20. Phylogenetic path model summaries for analyses restricted to species occupying the tropical biome, with population density as the response variable, across all combinations of predictor-response networks, trait evolution models and phylogenetic trees. Model statistics include the the number of independence claims made by the model (k), the number of model parameters (q), Fisher’s C goodness-of-fit statistic (C) and associated P-value (p), and the small-sample-corrected C information criterion (CICc).

Supplementary Table 21. Phylogenetic path model summaries for analyses restricted to species occupying the tropical biome, with effective population size as the response variable, across all combinations of predictor-response networks, trait evolution models and phylogenetic trees. Model statistics include the the number of independence claims made by the model (k), the number of model parameters (q), Fisher’s C goodness-of-fit statistic (C) and associated P-value (p), and the small-sample-corrected C information criterion (CICc).

Supplementary Table 22. Regression coefficient estimates for all predictor variables, including associated standard errors and P-values, for analyses including all species with population density as the response variable, across all supported models.

Supplementary Table 23. Regression coefficient estimates for all predictor variables, including associated standard errors and P-values, for analyses including all species with effective population size as the response variable, across all supported models.

Supplementary Table 24. Regression coefficient estimates for all predictor variables, including associated standard errors and P-values, for analyses restricted to Cetartiodactyla species with population density as the response variable, across all supported models.

Supplementary Table 25. Regression coefficient estimates for all predictor variables, including associated standard errors and P-values, for analyses restricted to Cetartiodactyla species with effective population size as the response variable, across all supported models.

Supplementary Table 26. Regression coefficient estimates for all predictor variables, including associated standard errors and P-values, for analyses restricted to Primates species with population density as the response variable, across all supported models.

Supplementary Table 27. Regression coefficient estimates for all predictor variables, including associated standard errors and P-values, for analyses restricted to Primates species with effective population size as the response variable, across all supported models.

Supplementary Table 28. Regression coefficient estimates for all predictor variables, including associated standard errors and P-values, for analyses restricted to Carnivora species with population density as the response variable, across all supported models.

Supplementary Table 29. Regression coefficient estimates for all predictor variables, including associated standard errors and P-values, for analyses restricted to Carnivora species with effective population size as the response variable, across all supported models.

Supplementary Table 30. Regression coefficient estimates for all predictor variables, including associated standard errors and P-values, for analyses restricted to herbivore species with population density as the response variable, across all supported models.

Supplementary Table 31. Regression coefficient estimates for all predictor variables, including associated standard errors and P-values, for analyses restricted to herbivore species with effective population size as the response variable, across all supported models.

Supplementary Table 32. Regression coefficient estimates for all predictor variables, including associated standard errors and P-values, for analyses restricted to omnivore species with population density as the response variable, across all supported models.

Supplementary Table 33. Regression coefficient estimates for all predictor variables, including associated standard errors and P-values, for analyses restricted to omnivore species with effective population size as the response variable, across all supported models.

Supplementary Table 34. Regression coefficient estimates for all predictor variables, including associated standard errors and P-values, for analyses restricted to carnivore species with population density as the response variable, across all supported models.

Supplementary Table 35. Regression coefficient estimates for all predictor variables, including associated standard errors and P-values, for analyses restricted to carnivore species with effective population size as the response variable, across all supported models.

Supplementary Table 36. Regression coefficient estimates for all predictor variables, including associated standard errors and P-values, for analyses restricted to species occupying the cold biome with population density as the response variable, across all supported models.

Supplementary Table 37. Regression coefficient estimates for all predictor variables, including associated standard errors and P-values, for analyses restricted to species occupying the cold biome with effective population size as the response variable, across all supported models.

Supplementary Table 38. Regression coefficient estimates for all predictor variables, including associated standard errors and P-values, for analyses restricted to species occupying the arid biome with population density as the response variable, across all supported models.

Supplementary Table 39. Regression coefficient estimates for all predictor variables, including associated standard errors and P-values, for analyses restricted to species occupying the arid biome with effective population size as the response variable, across all supported models.

Supplementary Table 40. Regression coefficient estimates for all predictor variables, including associated standard errors and P-values, for analyses restricted to species occupying the tropical biome with population density as the response variable, across all supported models.

Supplementary Table 41. Regression coefficient estimates for all predictor variables, including associated standard errors and P-values, for analyses restricted to species occupying the tropical biome with effective population size as the response variable, across all supported models.

Supplementary Table 42. Regression coefficient estimates for the adult mass variable, including associated standard errors and P-values, for EER-based analyses including all species with population density as the response variable, across all supported models.

Supplementary Table 43. Regression coefficient estimates for the adult mass variable, including associated standard errors and P-values, for EER-based analyses including all species with metabolic rate as the response variable, across all supported models.

Supplementary Table 44. Regression coefficient estimates for the adult mass variable, including associated standard errors and P-values, for EER-based analyses restricted to Cetartiodactyla species with population density as the response variable, across all supported models.

Supplementary Table 45. Regression coefficient estimates for the adult mass variable, including associated standard errors and P-values, for EER-based analyses restricted to Cetartiodactyla species with metabolic rate as the response variable, across all supported models.

Supplementary Table 46. Regression coefficient estimates for the adult mass variable, including associated standard errors and P-values, for EER-based analyses restricted to Primates species with population density as the response variable, across all supported models.

Supplementary Table 47. Regression coefficient estimates for the adult mass variable, including associated standard errors and P-values, for EER-based analyses restricted to Primates species with metabolic rate as the response variable, across all supported models.

Supplementary Table 48. Regression coefficient estimates for the adult mass variable, including associated standard errors and P-values, for EER-based analyses restricted to Carnivora species with population density as the response variable, across all supported models.

Supplementary Table 49. Regression coefficient estimates for the adult mass variable, including associated standard errors and P-values, for EER-based analyses restricted to Carnivora species with metabolic rate as the response variable, across all supported models.

Supplementary Table 50. Distributions of supported models with respect to the underlying predictor-response networks, for all combinations of data subsets and response variables.

