## Supplementary figures and images for "Trade-offs beget trade-offs: Causal analysis of mammalian population dynamics"

### supp_fig_1.pdf

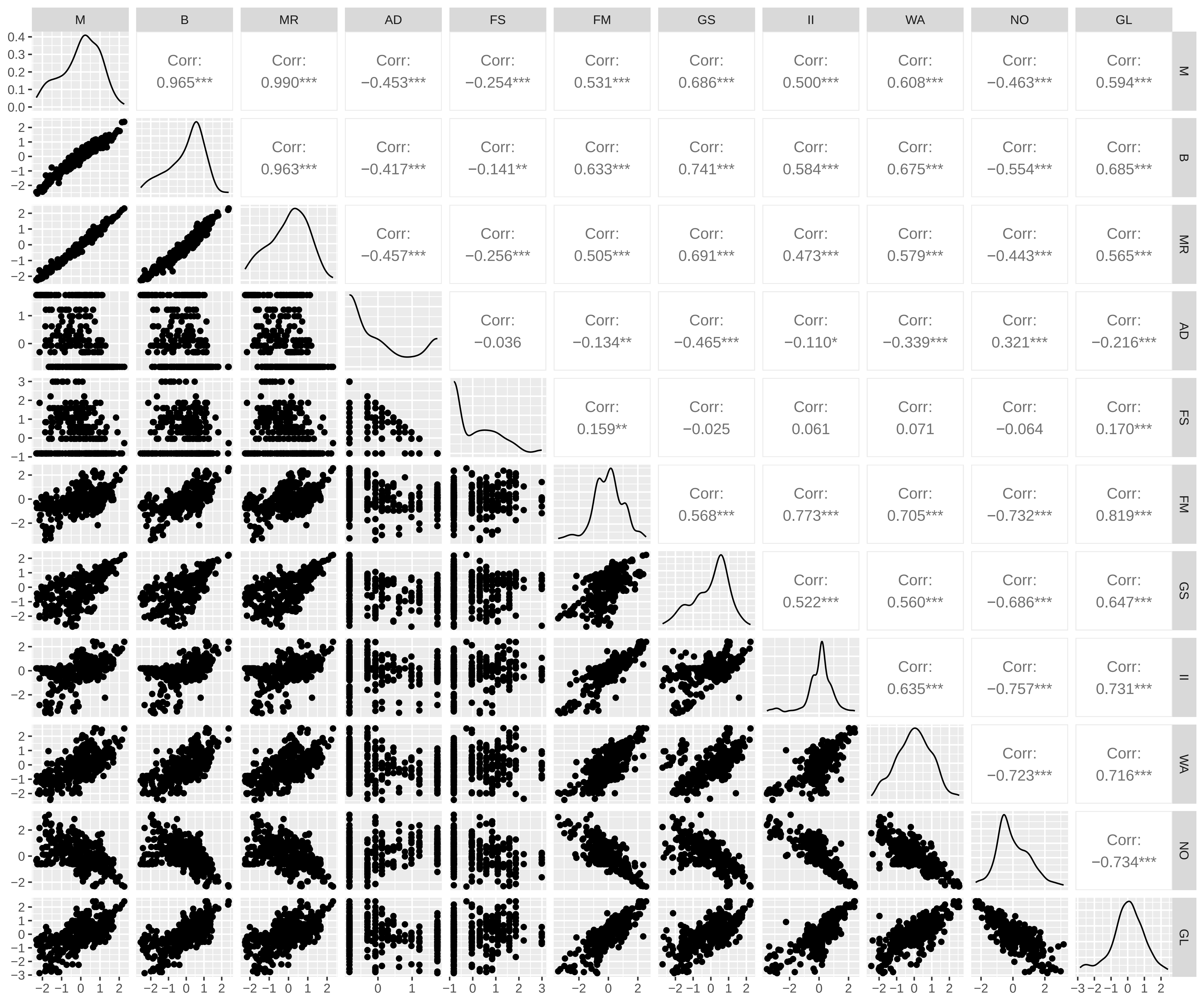

### supp_fig_2.pdf

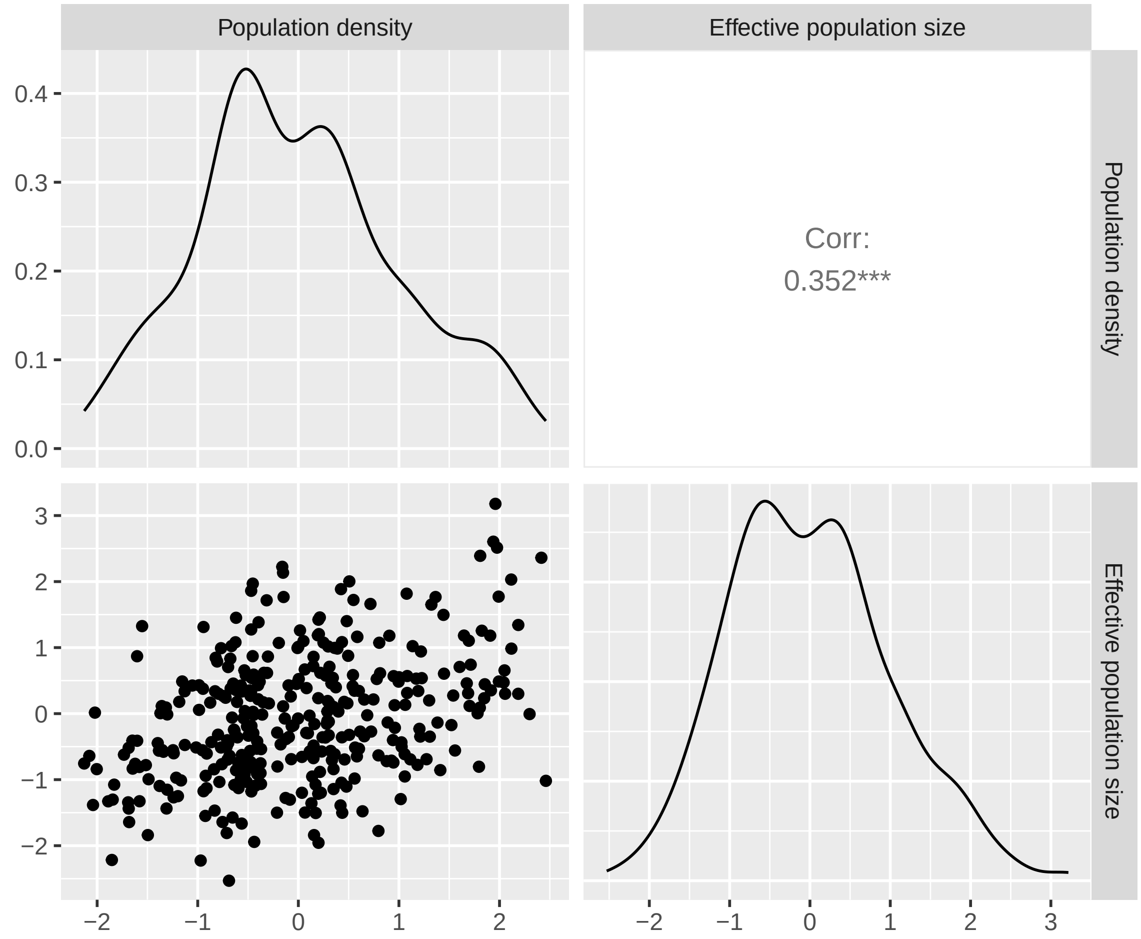

### supp_fig_3.pdf

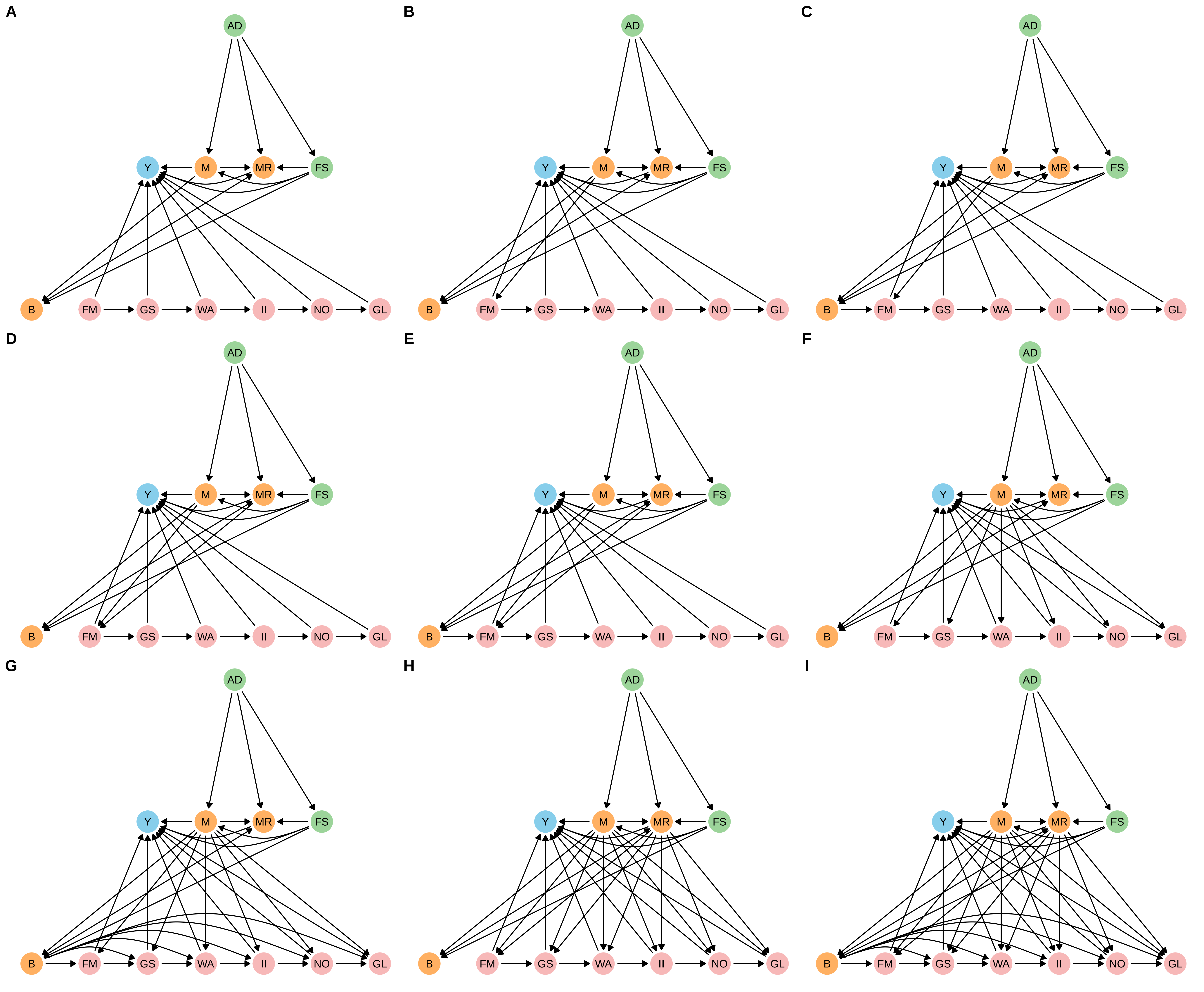

### supp_fig_4.pdf

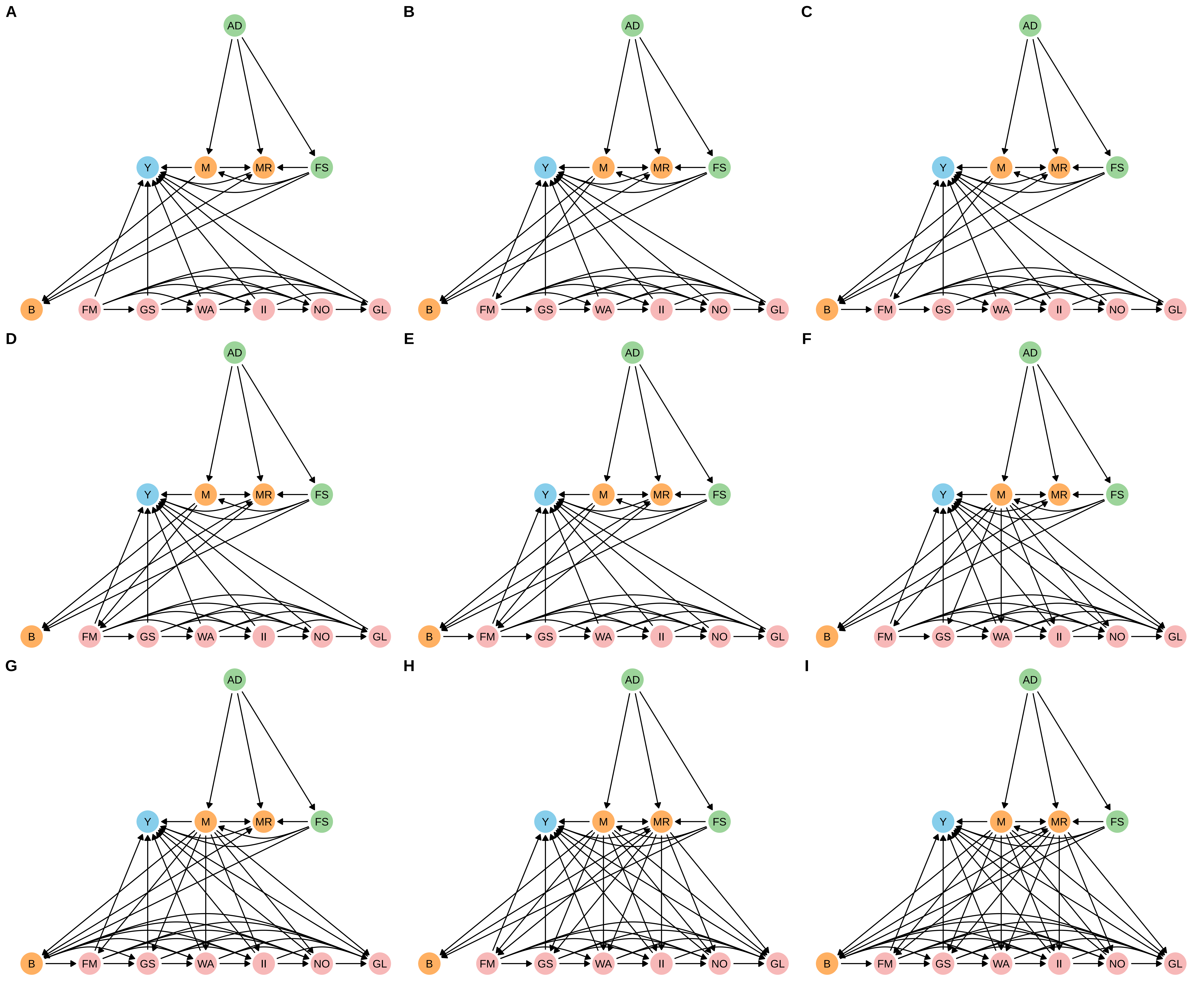

### supp_fig_5.pdf

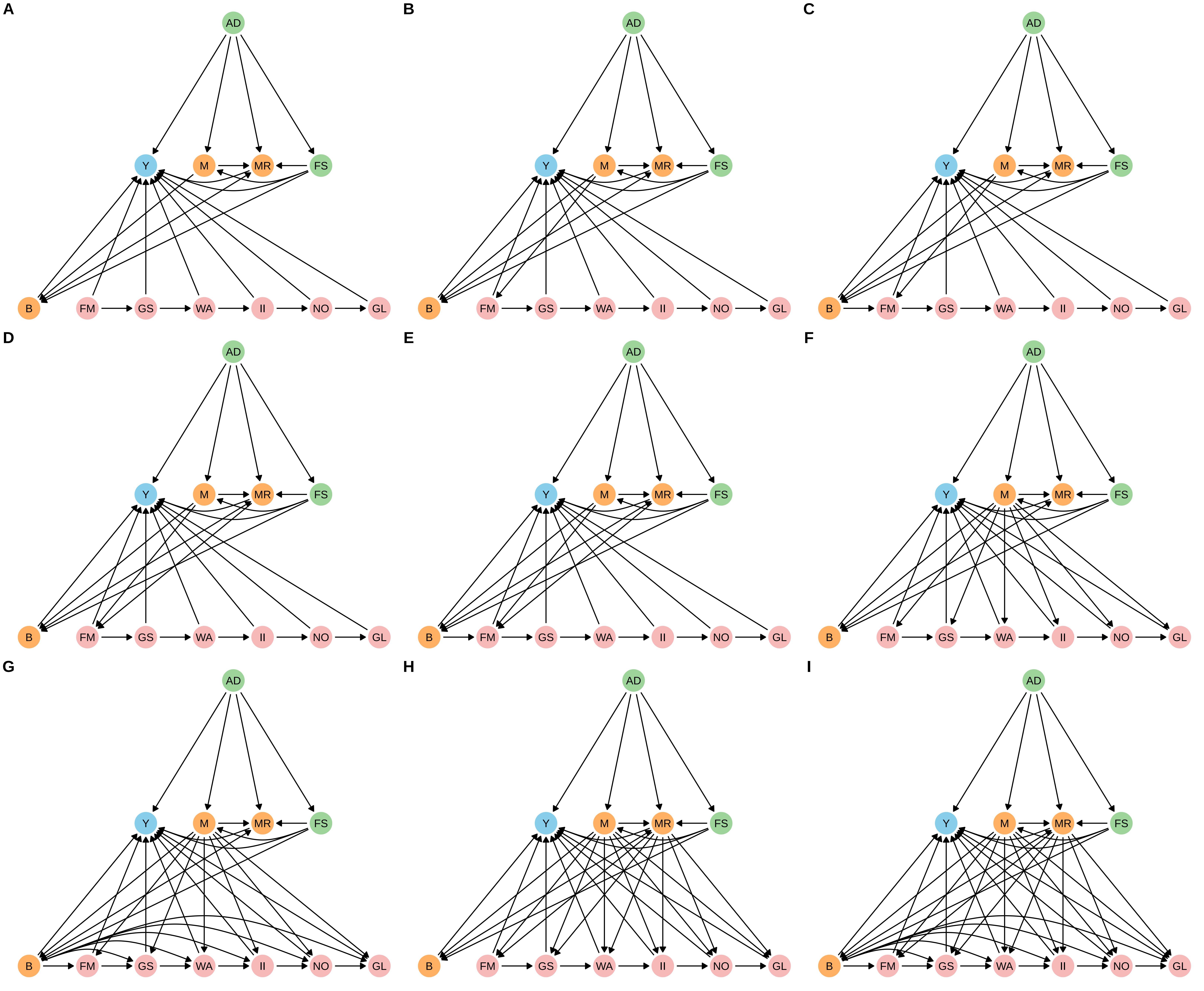

### supp_fig_6.pdf

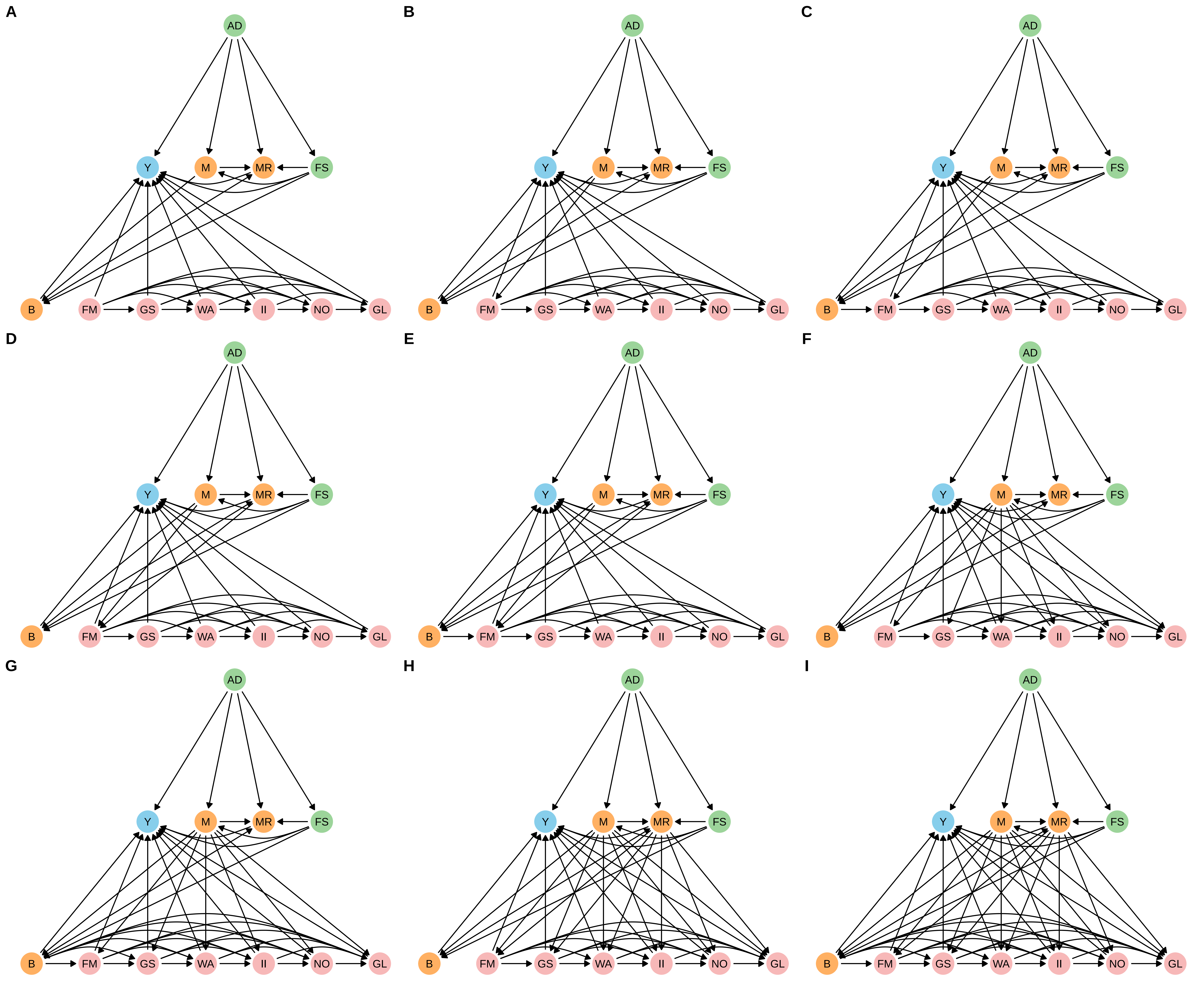

### supp_fig_7.pdf

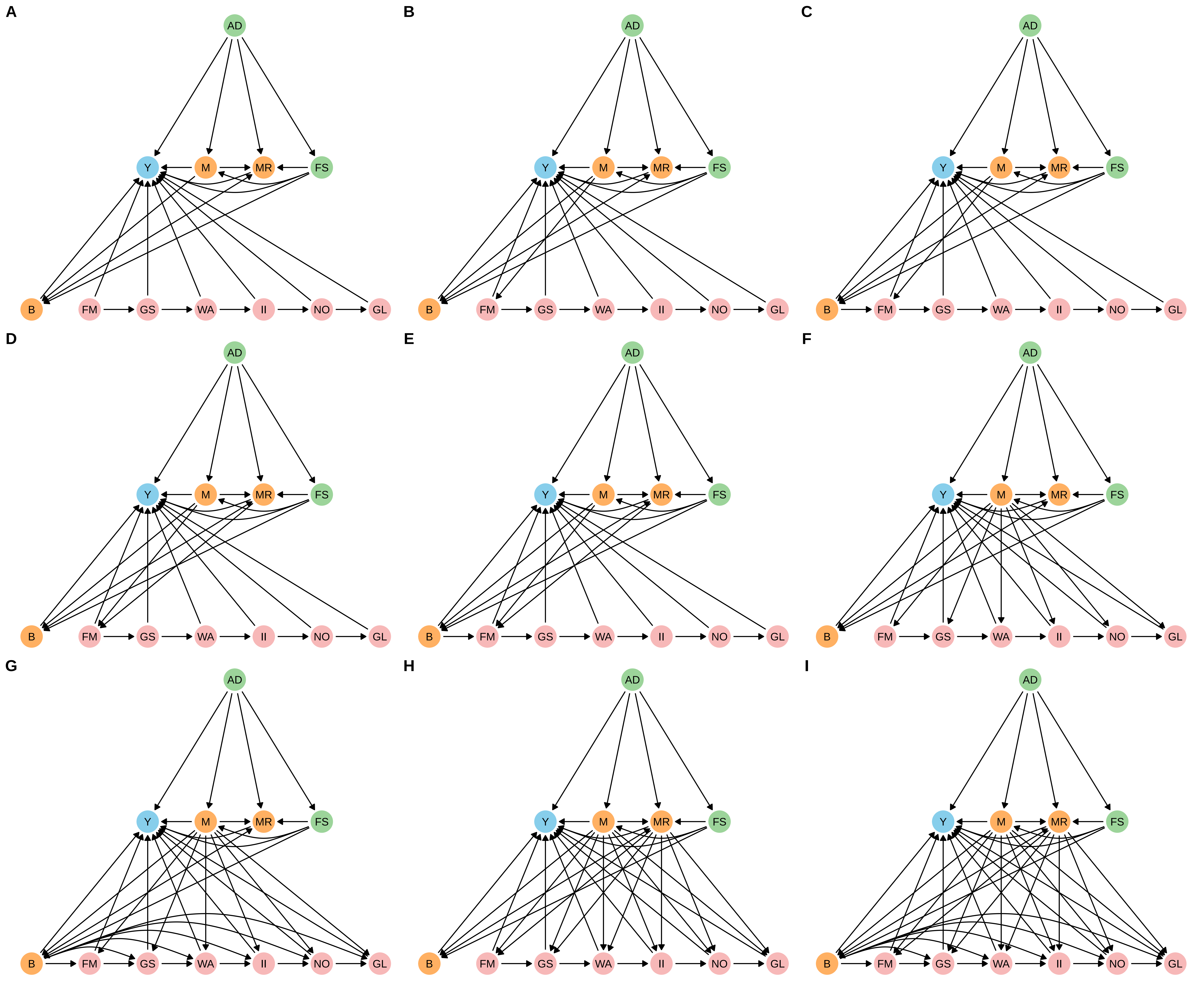

### supp_fig_8.pdf

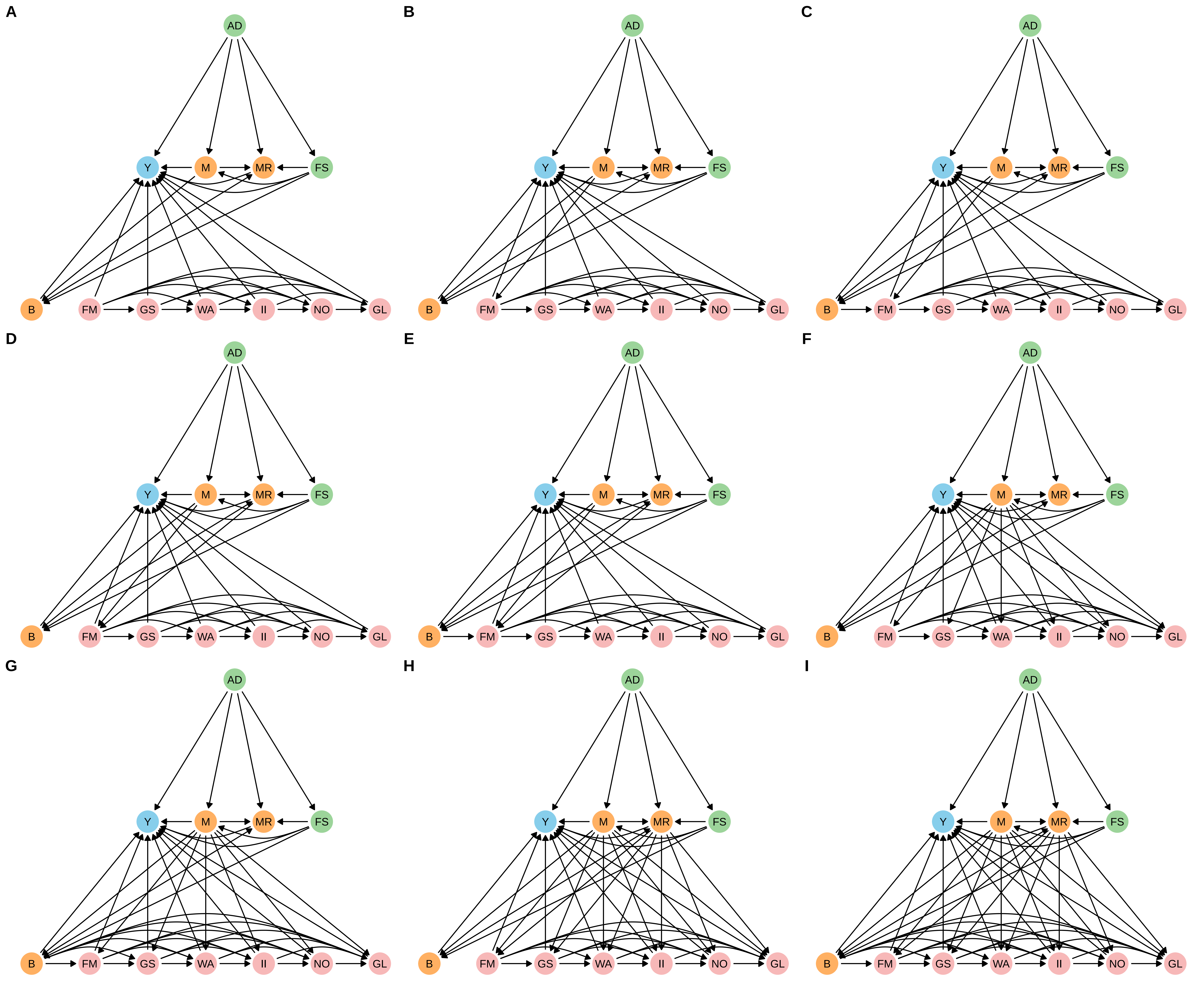

### supp_fig_9.pdf

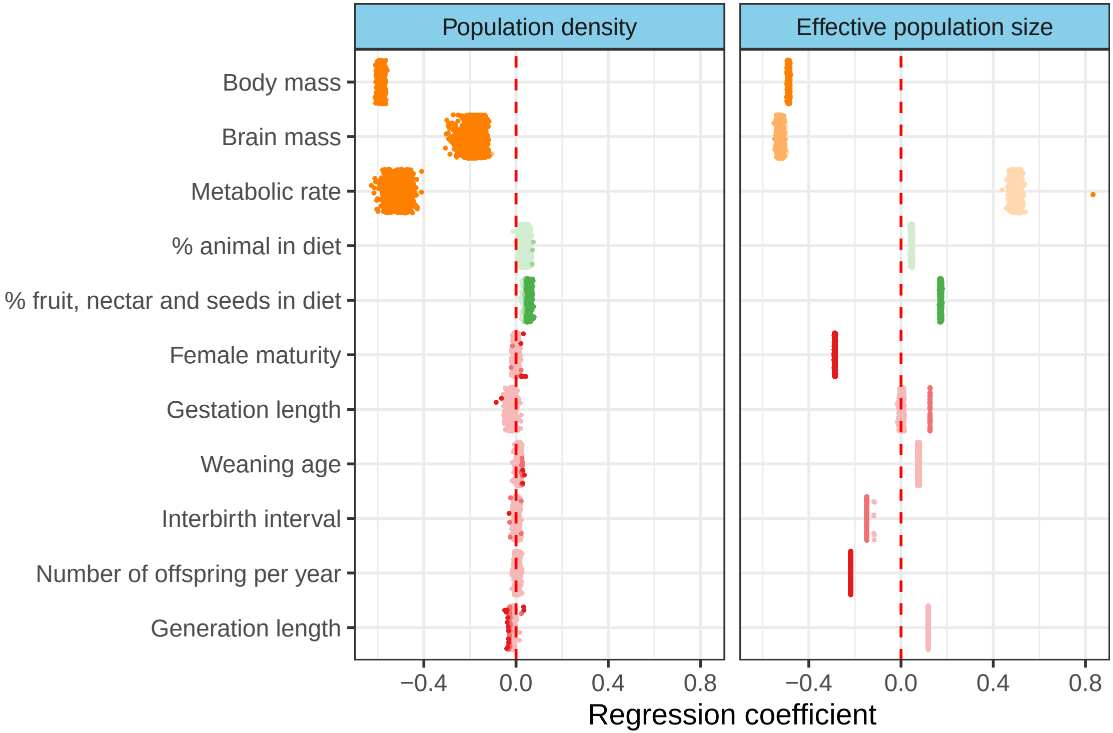

### supp_fig_10.pdf

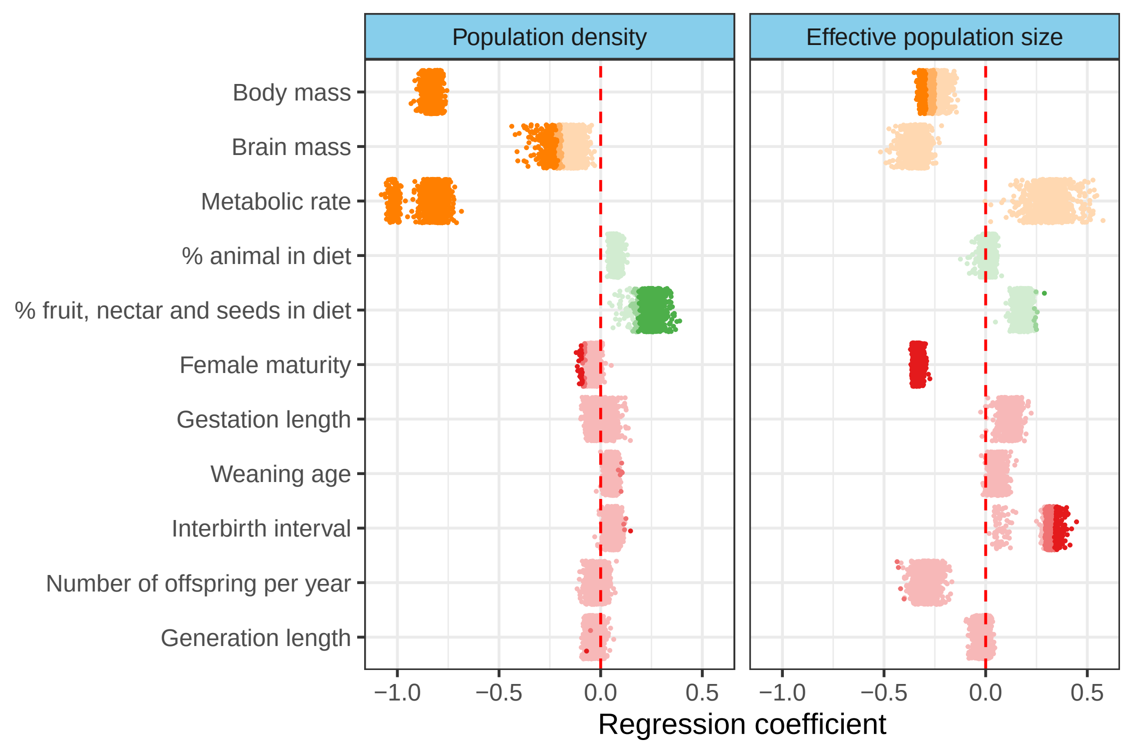

### supp_fig_11.pdf

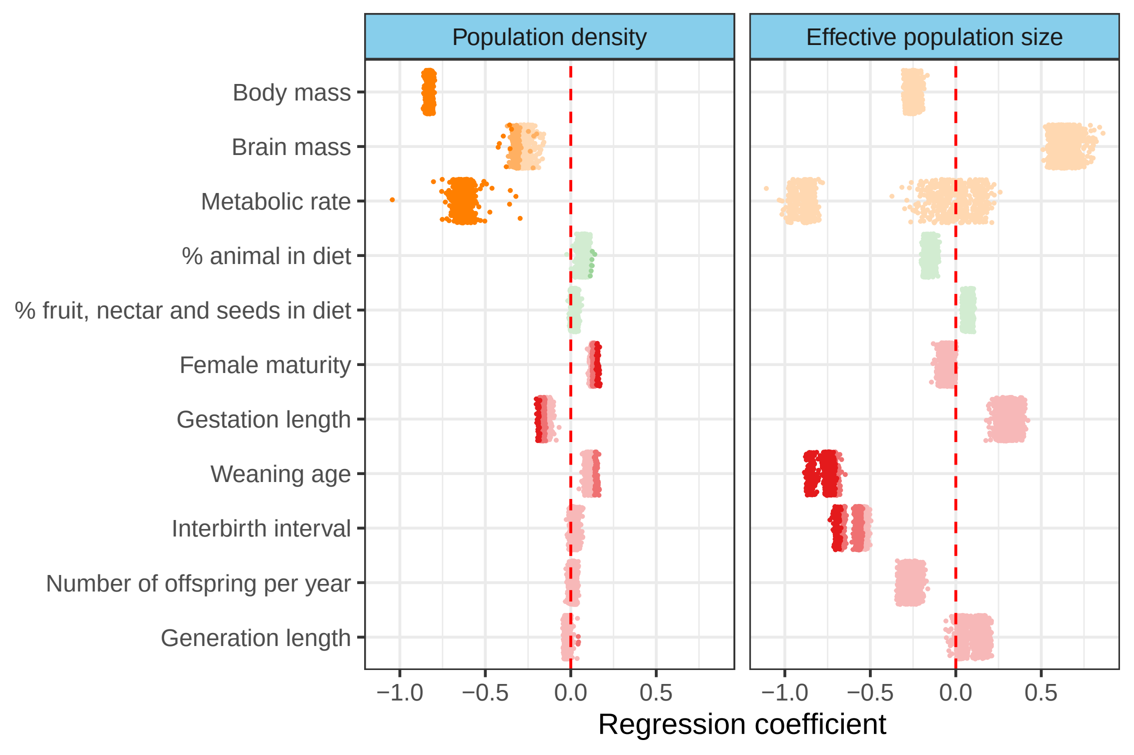

### supp_fig_12.pdf

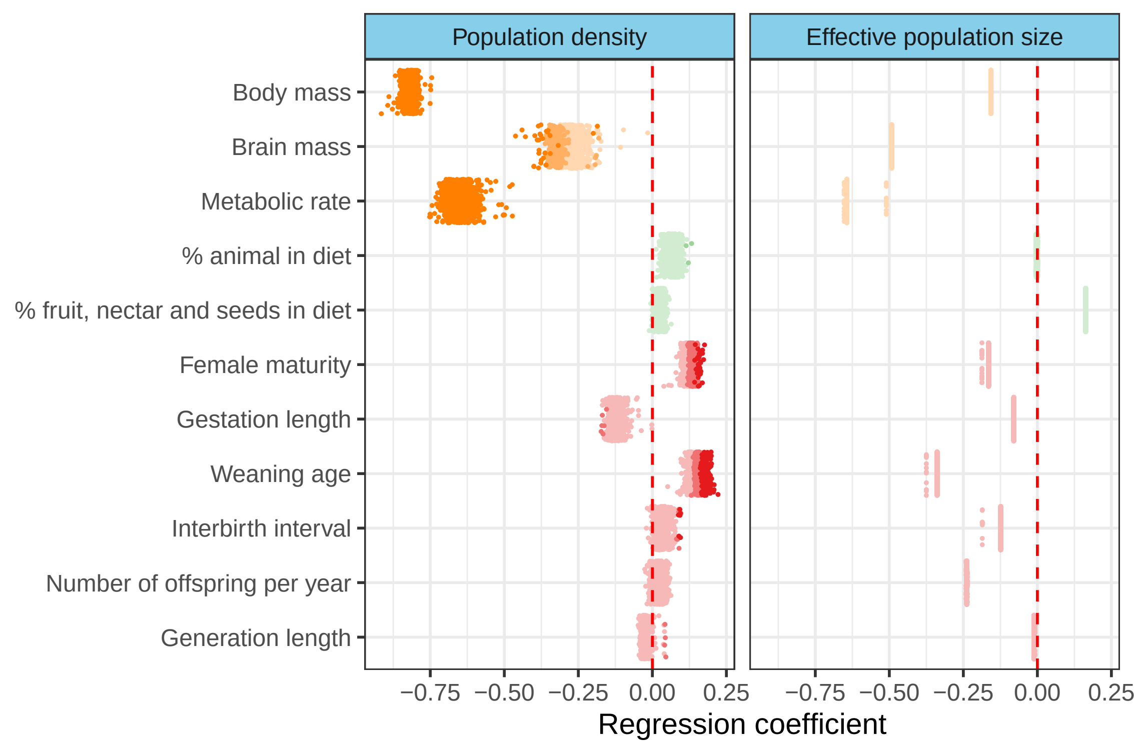

### supp_fig_13.pdf

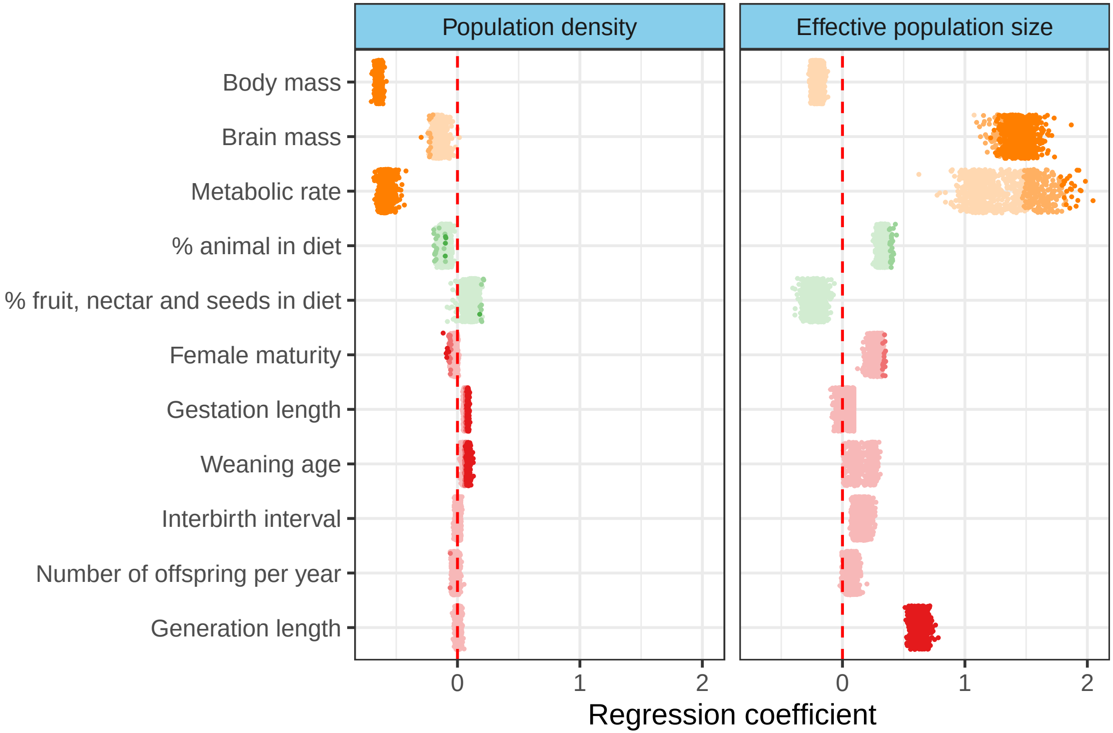

### supp_fig_14.pdf

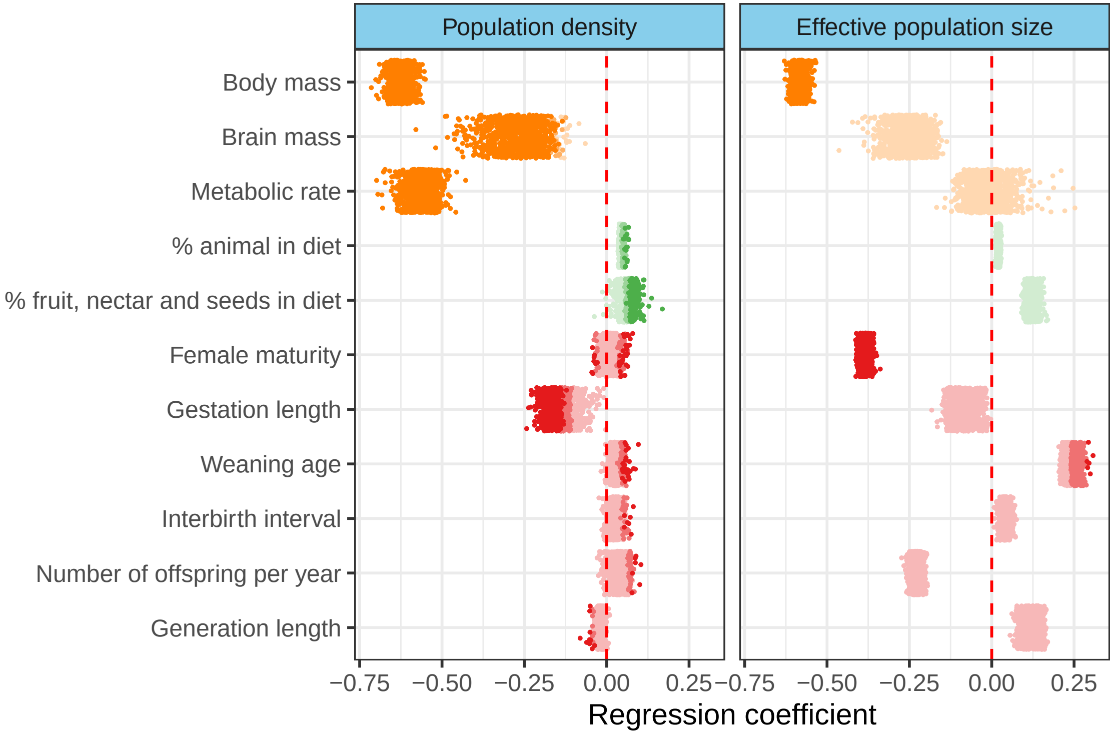

### supp_fig_15.pdf

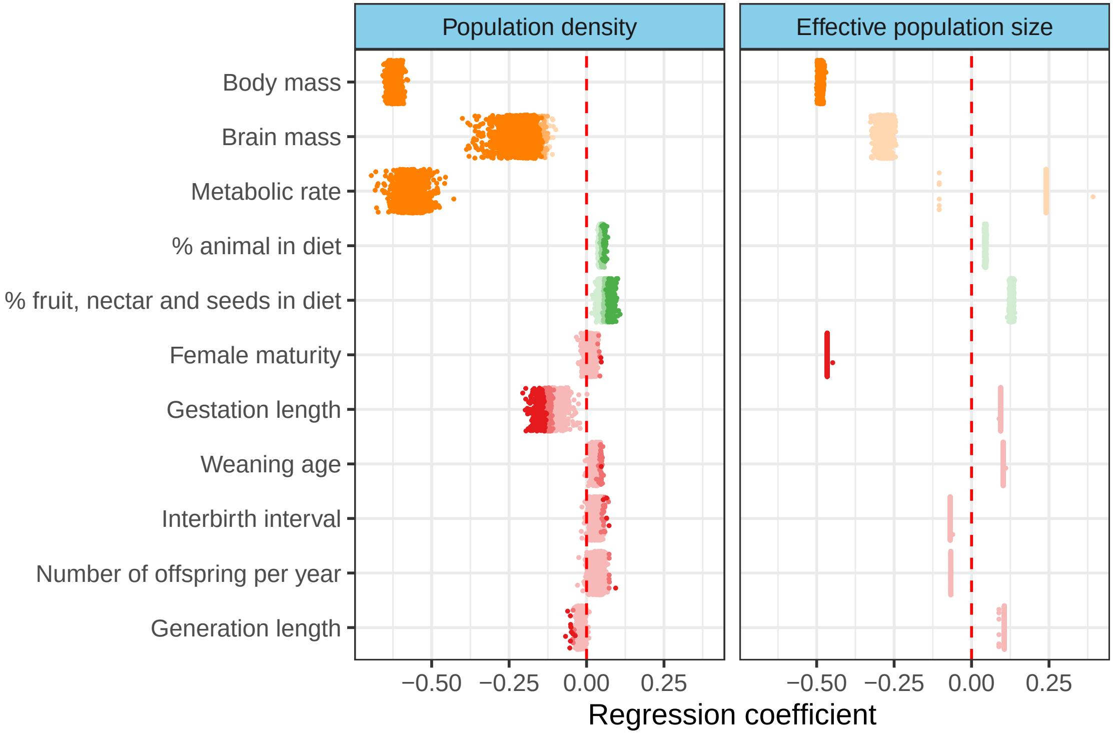

### supp_fig_16.pdf

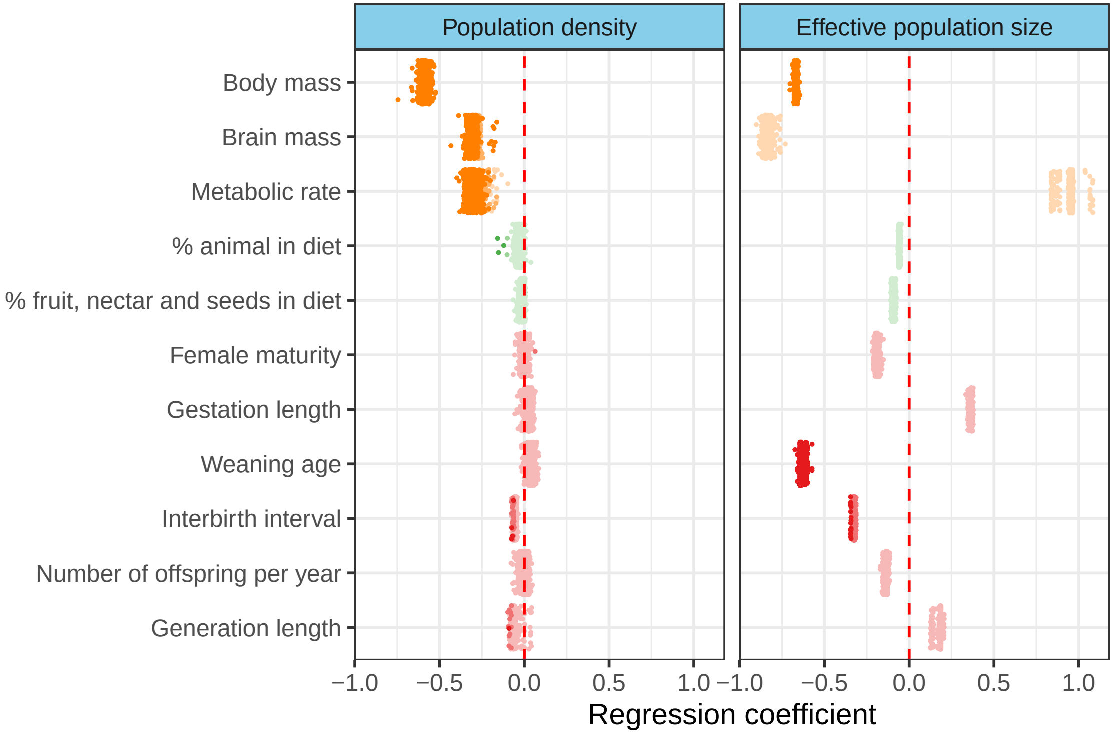

### supp_fig_17.pdf

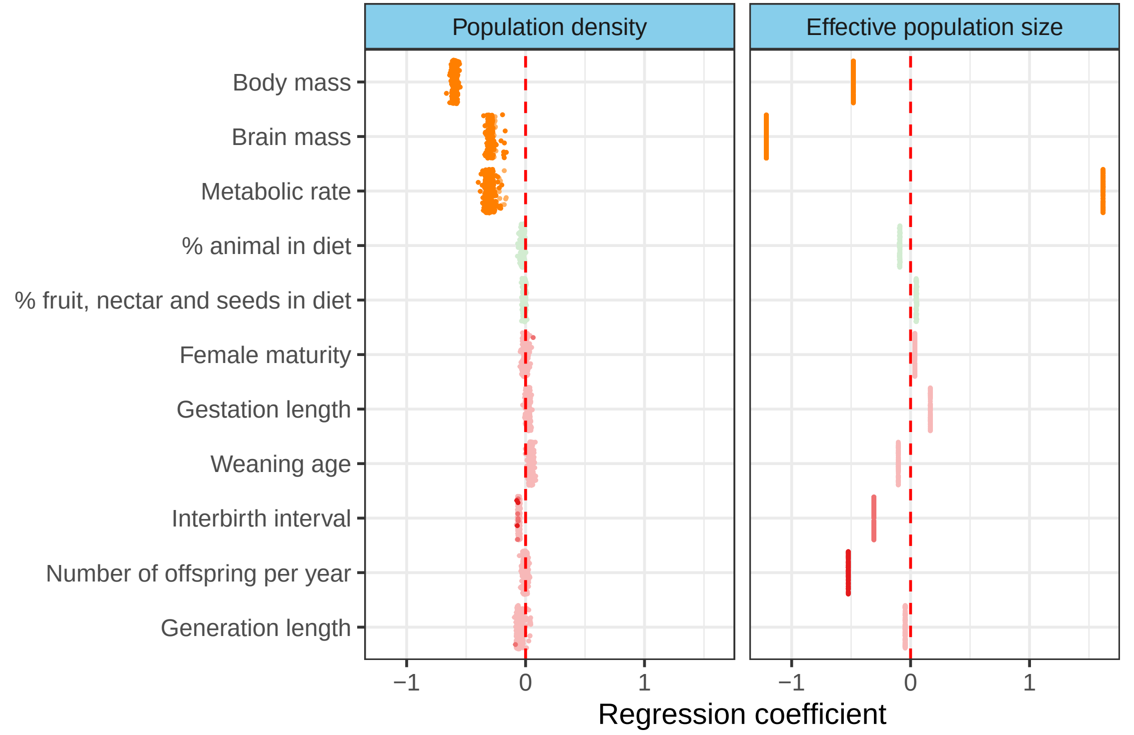

### supp_fig_18.pdf

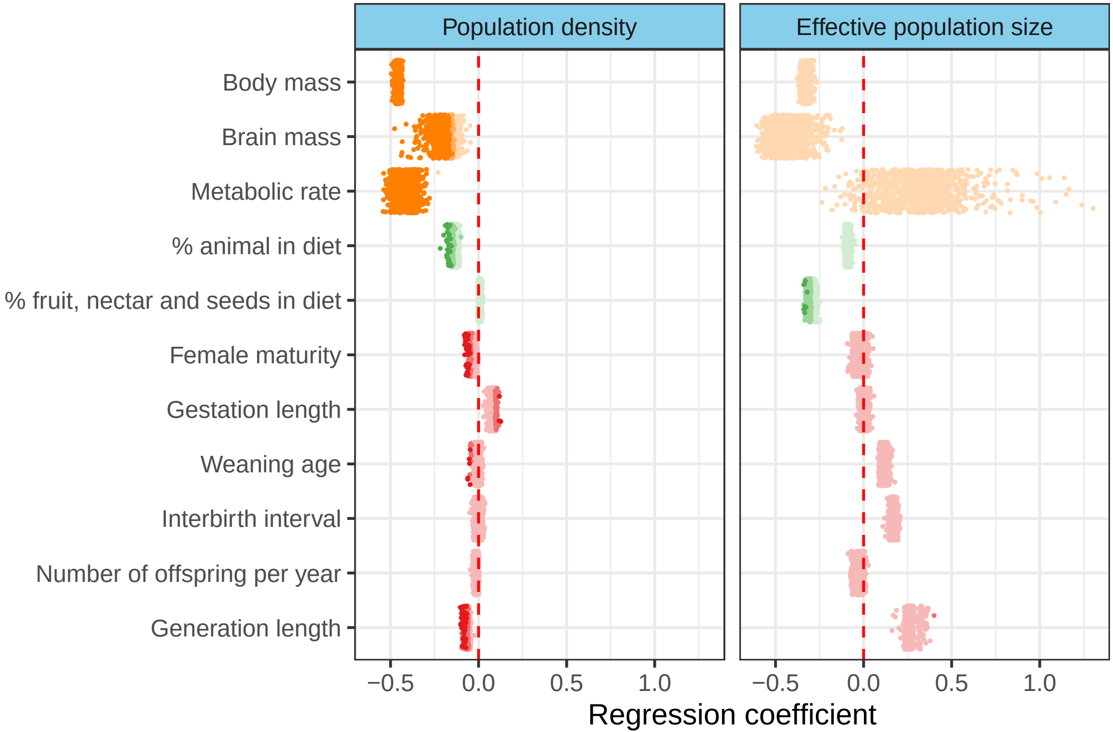

### supp_fig_19.pdf

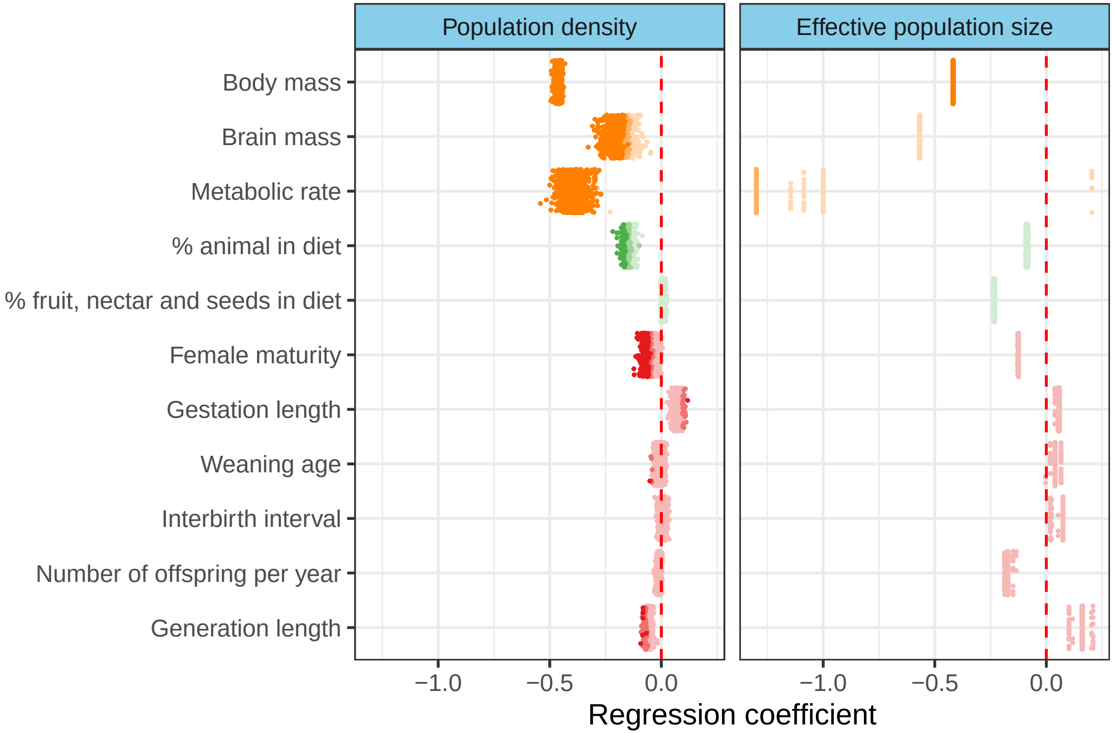

### supp_fig_20.pdf

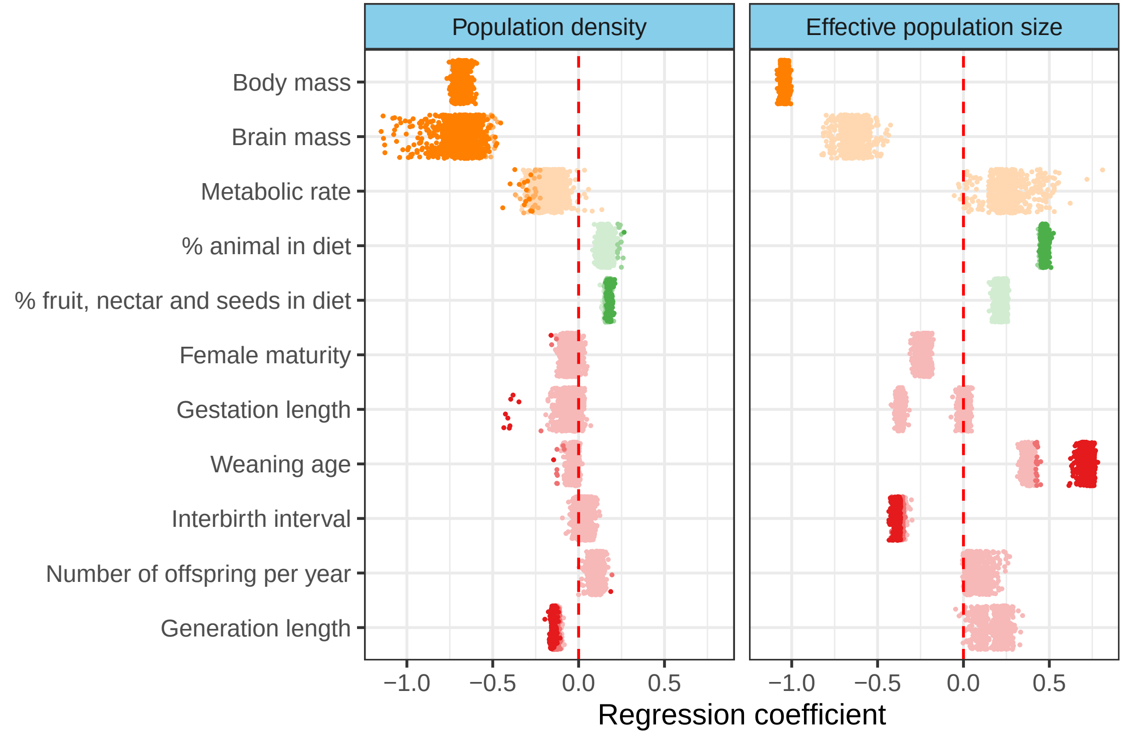

### supp_fig_21.pdf

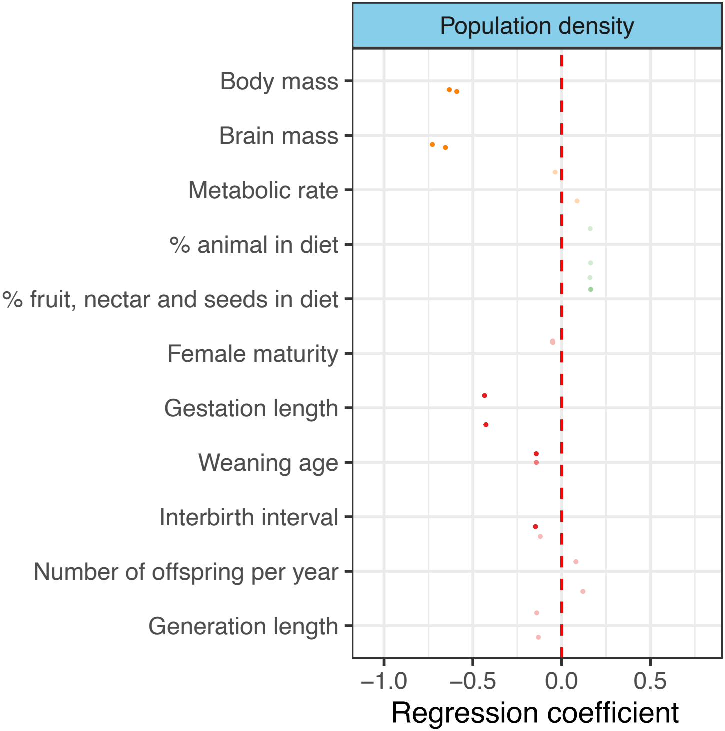

### supp_fig_22.pdf

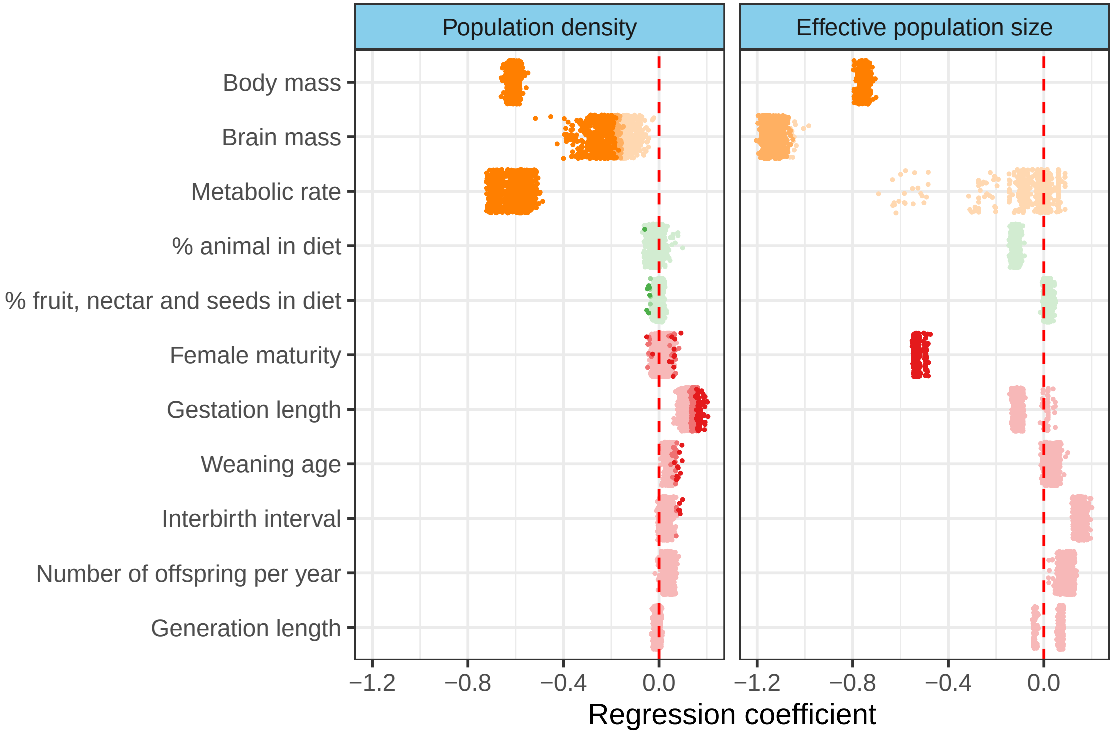

### supp_fig_23.pdf

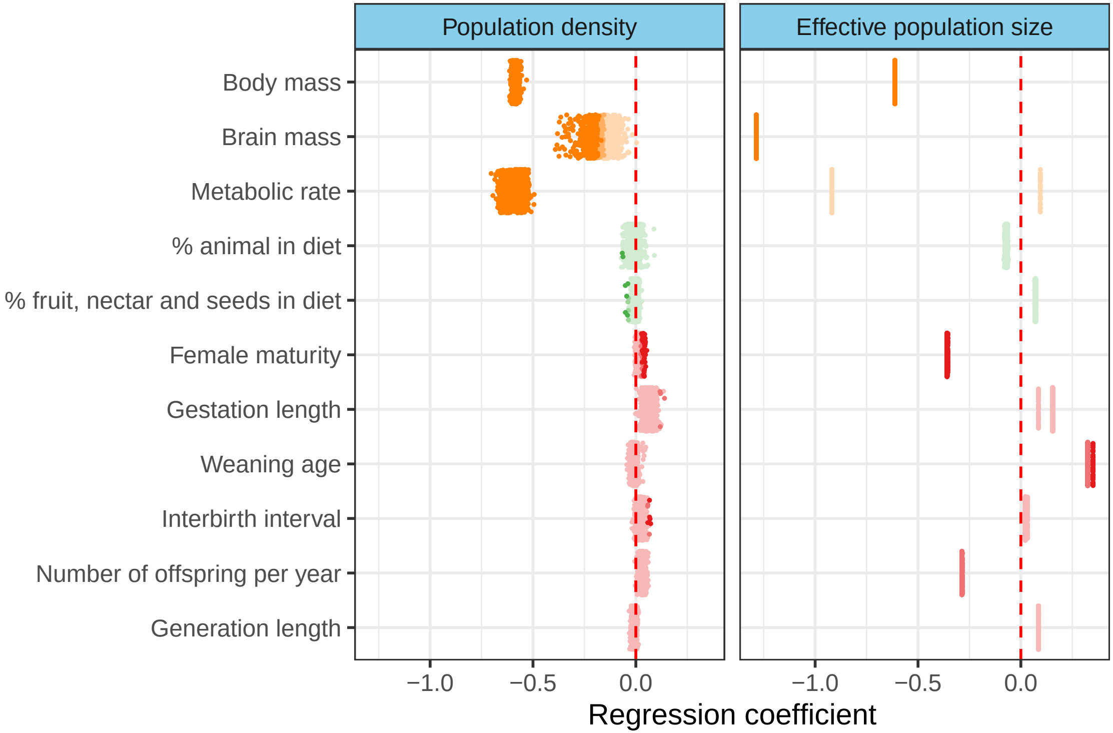

### supp_fig_24.pdf

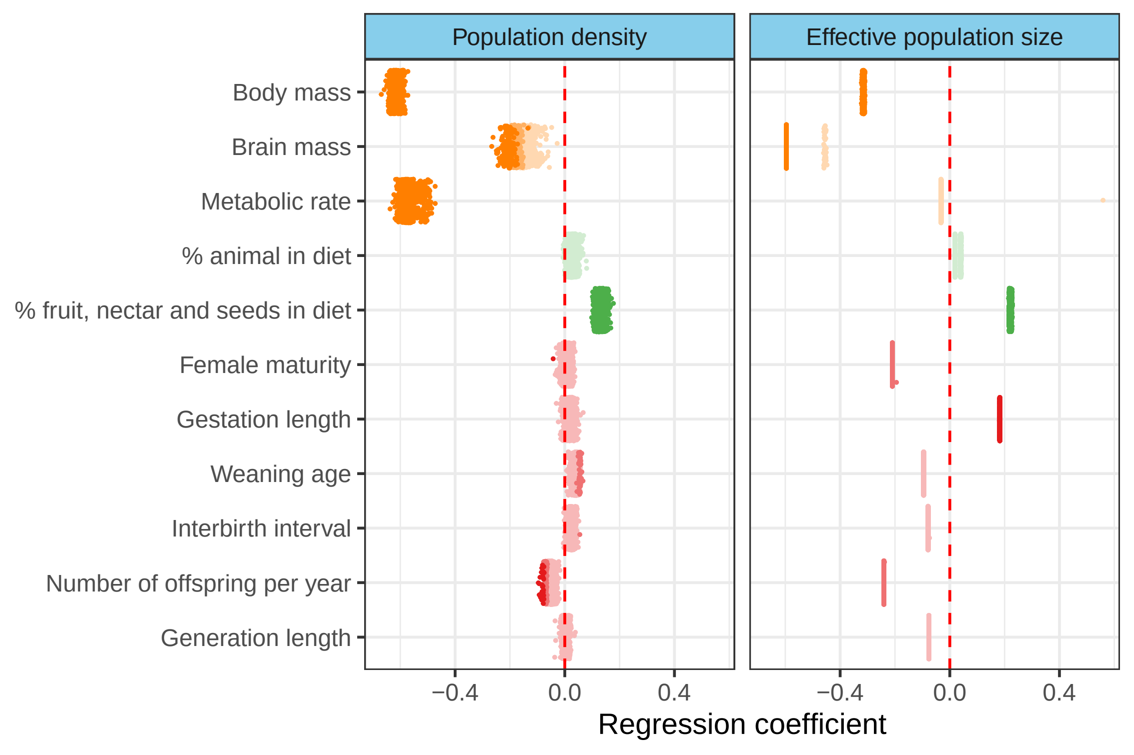

### supp_fig_25.pdf

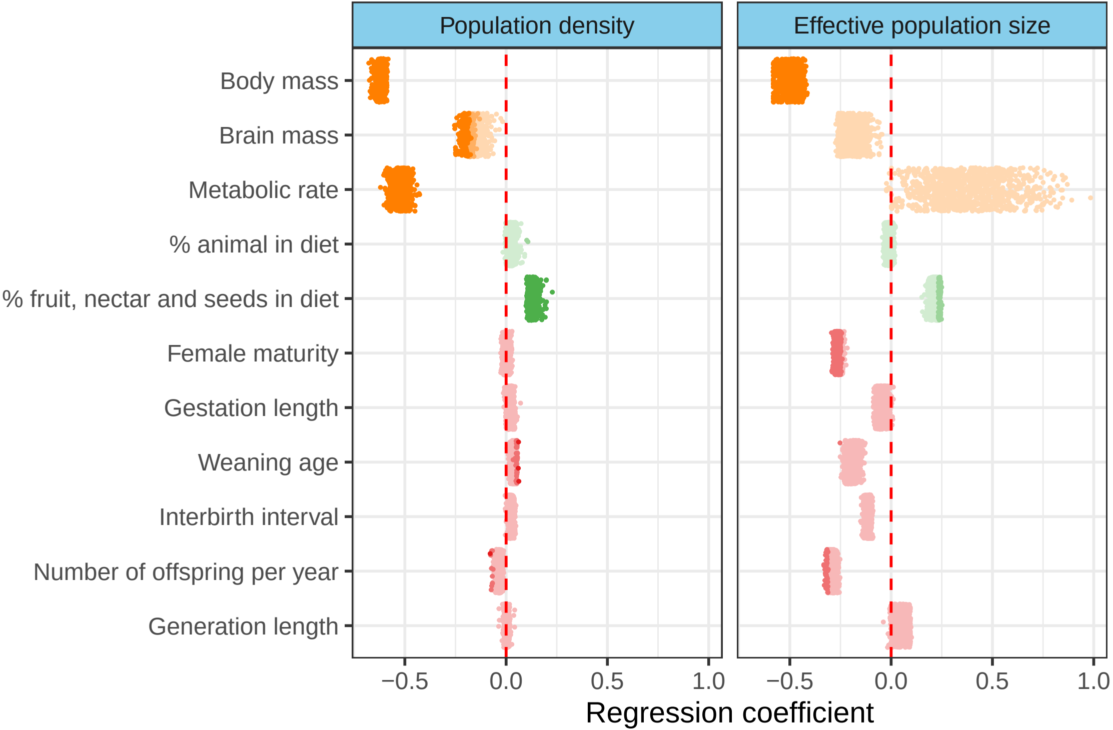
